# A role of extracellular vesicle-mediated inter-organ communication in obesity-related arrhythmia

**DOI:** 10.1101/2025.09.13.676027

**Authors:** Worawan B. Limpitikul, Marta Garcia-Contreras, Michael J. Betti, Paul Spangler, Quanhu Sheng, Steffen Pabel, Samuel Sossalla, Ling Xiao, Emeli Chatterjee, Patrick T. Ellinor, Eric R. Gamazon, Ravi Shah, Saumya Das

## Abstract

Obesity contributes to the risk of cardiac arrhythmias, but the exact mechanism remains unclear. Here, we show visceral adipose tissue-derived extracellular vesicles (VAT EVs) from individuals with obesity prolong action potential duration (APD) and impair calcium handling in stem cell-derived cardiomyocytes, in addition to activating fibroblasts and macrophages towards a pro-fibrotic/inflammatory state, thereby creating pro-arrhythmic substrate. Adipose-derived EVs target the heart in obese mice, suggesting the potential for direct communication. Transcriptome-wide genetic association (TWAS) and epigenetic studies anchored on genes differentially expressed in cardiomyocytes, fibroblasts, and macrophages after VAT-EV exposure identified genes significantly associated with QT interval and atrial fibrillation. Finally, as a proof-of-principle, we pharmacologically blocked TRPC3 (a VAT-EV-induced ion channel) in cardiomyocytes, restoring the APD towards normality. This molecular genetic evidence supports an EV-mediated direct communication pathway between adipose tissue and the heart in arrhythmogenesis, offering a new paradigm to identify mediators of cardiovascular disease in obesity.

**Graphical Abstract:** 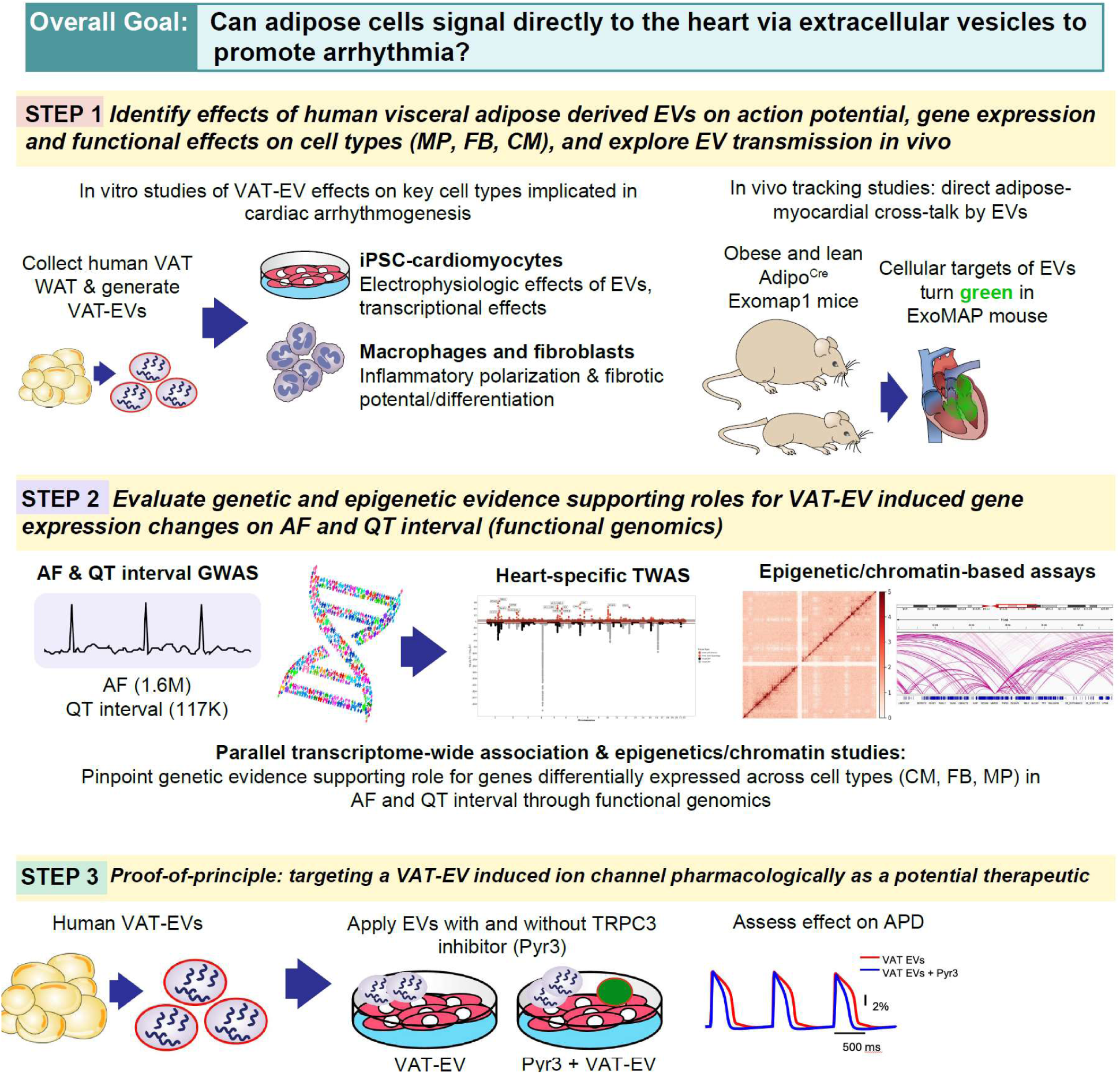

## INTRODUCTION

With nearly 2.5 billion overweight or obese^1^, obesity exacts an increasing, profound toll on global cardiometabolic health. Alongside the metabolic complications accompanying increasing body mass index (BMI), obesity drives development and progression of atrial fibrillation (AF)^2–6^, and obesity-related AF appears more resistant to conventional therapeutic interventions (e.g., catheter ablation^7,8^ and antiarrhythmic therapy^9^). Moreover, each 5-unit increase in BMI portends a 16% increase hazard of sudden cardiac death^10^, and individuals with obesity harbor markers of increased susceptibility to ventricular arrhythmias, including ventricular ectopy^11^ and prolonged QT interval^12^. In effect, reductions in obesity or obesity-related cardiometabolic risk have been associated with improvements in AF^13–15^, and recent genetic studies pinpoint a shared causal predisposition between obesity and AF^16^. However, what confers this predisposition remains occult: while obesity is associated with a host of <u>systemic</u> and <u>myocardial</u> states that predispose to atrial and ventricular arrhythmia (e.g., inflammation, mitochondrial dysfunction, ventricular-vascular coupling^17–19^), studies in vascular biology suggest that local paracrine or endocrine factors may signal between fat and myocardium to increase susceptibility to cardiovascular disease^20^. Recent work in our group has demonstrated potential for extracellular vesicles (EVs)—membrane-bound vesicles that can carry molecular signals (e.g., RNAs, proteins) across tissues—in the pathogenesis of obesity-related complications^21^. However, the role for EVs as a novel paradigm for fat-myocardial communication in obesity-related AF has not been demonstrated.

Here, we hypothesize direct communication between visceral adipose tissue (VAT) and myocardium causally contributes to obesity-related cardiac arrhythmogenesis. For the first time to our knowledge, we demonstrate that atrial myocytes from human atrial tissue exhibit a <u>prolonged</u> action potential duration (APD) in individuals with obesity, in contrast to widely used animal AF models with shortened APD^22^. We subsequently show that EVs isolated from human VAT from individuals with obesity produced three key pathophysiological changes in cardiac cells: (1) significant APD prolongation in both atrial and ventricular induced pluripotent stem cell-derived cardiomyocytes; (2) a profibrotic phenotype in cardiac fibroblasts, with enhanced myofibroblast differentiation and increased collagen production; and (3) pro-inflammatory polarization in cardiac macrophages linked to arrhythmogenesis^23–25^. Using a genetic model of EV tracking (*Exomap1*)^26^ to trace adipose-derived EVs, we demonstrate *in vivo* evidence of adipose-derived EV delivery to the myocardium, specifically in obese mice. Next, to identify potential causal mechanisms of visceral adipose tissue-derived extracellular vesicles (VAT EVs) on arrhythmogenesis, we performed transcriptomics in VAT EV-treated iPSC-CMs, demonstrating (1) enrichment for pathways putatively involved in arrhythmogenesis and (2) concordant causal mapping to large human genome-wide association studies (GWAS) of AF and QT prolongation. We experimentally validated a prioritized target—*TRPC3*, a nonselective cation channel—targeted by VAT EV signaling, with TRPC3 inhibition abrogating VAT EV effects on APD prolongation in iPSC-CMs. Collectively, these findings establish adipose-derived EV transmission as a novel avenue of interorgan crosstalk in obesity-related arrhythmias, expanding the current mechanistic paradigm linking excess adiposity to cardiovascular risk in obesity.

## RESULTS

### Altered electrical properties of myocytes in obese individuals

To explore the effect of obesity on electrophysiological properties of atrial myocytes, we first examined the relationship between obesity and cardiac electrophysiology in human subjects. Action potential duration at 90% repolarization (APD_90_) was measured at 1 Hz pacing rate in cardiac tissue samples from patients undergoing open heart surgeries with BMI <25 kg/m^2^ (lean) and BMI >25 kg/m^2^ (overweight/obese). **Fig. 1a** demonstrates action potential tracing from myocytes isolated from right atria of individuals with BMI < 25 and > 25 kg/m^2^. Patients with higher BMI exhibit significantly prolonged action potential duration compared to lean individuals (**Fig. 1b**, 157.5±71.4ms vs 77.9±28.4 ms, p = 0.003), providing clinical evidence that obesity is associated with altered cardiac electrophysiology that could predispose to arrhythmias. Detailed characteristics of right atrial tissue donors are displayed in **Supplementary Table 1**.

**Figure 1:**
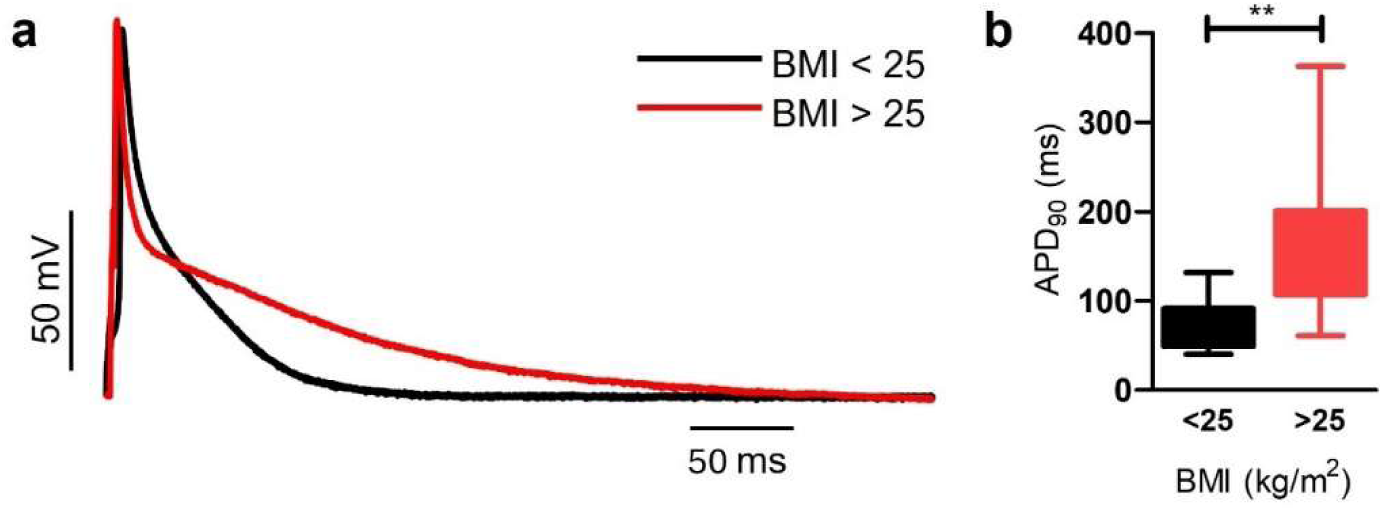
Altered electrical properties of myocytes in obese individuals. a, Action potential in human right atrial myocytes isolated from patients undergoing open heart surgeries. **b,** Action potential duration measured at 90% repolarization (APD₉₀). Atrial myocytes from lean patients (BMI < 25 kg/m^2^; n=9 cells) showed significantly shorter APD₉₀ compared to atrial myocytes from overweight/obese patients (BMI > 25 kg/m^2^; n=27 cells) measured at 1 Hz pacing rate. Individual data points represent measurements from single myocytes, with bars showing mean±SEM.

### VAT EVs promote pro-arrhythmic electrical and Ca^2+^ activity in iPSC-derived cardiomyocytes

We next treated iPSC-derived atrial cardiomyocytes (iPSC-aCMs) and iPSC-derived ventricular cardiomyocytes (iPSC-vCMs) with EVs derived from VAT to assess electrical properties. EVs were extracted from conditioned medium of cultured minced VAT (VAT EVs) from donors who underwent bariatric surgeries (NCT06408961) by size exclusion chromatography. Detailed characteristics of VAT donors are displayed in **Supplementary Table 2**. VAT EVs as well as EVs derived from 0.5 ml of plasma from obese (BMI > 30 kg/m^2^) or lean (BMI < 25 kg/m^2^) patients were characterized following MISEV (Minimal Information for Studies of Extracellular Vesicles), including size distribution and concentration via microfluidic resistive pulse sensing (MRPS) and canonical surface and cargo protein markers with some variability observed in the yield of VAT-EVs obtained from the adipose tissue samples (**Supplementary Fig. 1**). Additional VAT EV characterization can be found in Chatterjee et al^21^. Both iPSC-aCMs and iPSC-vCMs were treated for 24 hours with VAT EVs at a concentration of 7.5 x 10^10^ EVs/mL. To account for potential unintended effects of EV treatment, the control group was treated with the same number of EVs extracted from condition media of iPSC-aCMs or iPSC-vCMs. Dose-response studies showed that treatment with of 7.5 x 10^10^ EVs/mL (but not lower doses) of VAT EVs for 24 hours rendered phenotypic changes without overt toxicity in iPSC-CMs (**Supplementary Fig. 2**)

**Fig. 2a** shows fluorescence signal representing membrane voltage of iPSC-aCMs during 1 Hz pacing. After 24 hours of VAT EV treatment, pronounced action potential prolongation was observed in iPSC-aCMs treated with VAT EVs compared to control. APD measured at 80% repolarization (*APD_80_*) at 1 Hz pacing rate was significantly longer in VAT EV-treated iPSC-aCMs compared to control (**Fig. 2b**, 394±28 ms vs 345±29 ms, p = 0.0002). Similarly, prolonged action potential was observed in iPSC-vCMs treated with VAT EVs (**Fig. 2c**) with significantly longer *APD_80_* compared to control (**Fig. 2d**, 602±56 ms vs 435±42 ms, p = 0.0001). Although VAT EV treatment led to prolongation of APD in both atrial and ventricular myocytes, an argument can be made that in this setting, cardiomyocytes are directly exposed to adipose tissue-derived EVs and may not represent the *in vivo* state in which cardiomyocytes are exposed to circulating EVs in which only a fraction of total EVs originate from adipose tissue, thus potentially diluting out the phenotype observed. To simulate this *in vivo* condition, we treated cardiomyocytes with plasma-derived EVs from lean and obese donors (see **Supplementary Table 3** for detailed characteristics of plasma donors). After 24-hour treatment of plasma-derived EVs, the action potential was similarly prolonged in iPSC-aCMs treated with plasma-derived EVs from obese donors compared to those from lean donors (**Supplementary Fig. 3a-b**). Similarly, iPSC-vCMs treated with plasma-derived EV from obese donors showed APD prolongation compared to those treated with EVs from lean donors (**Supplementary Fig. 3c-d**). In both cell types, the extent of APD prolongation after plasma EV treatment was less than after VAT EV treatment, likely reflecting exposure to relatively lower dose of adipose EVs in the plasma (compared to EVs directly derived from adipose explants).

**Figure 2:**
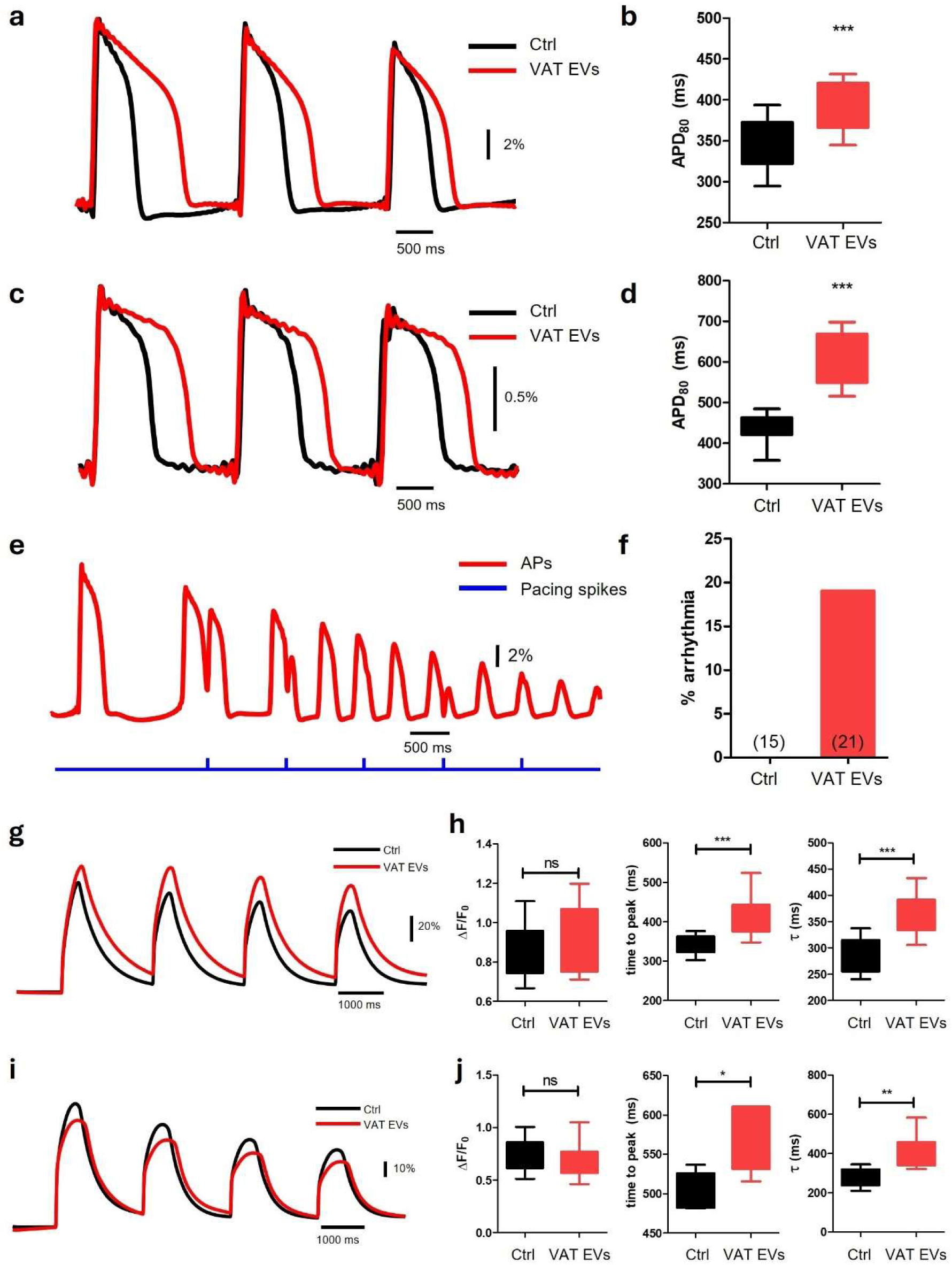
Alterations in electrical and calcium cycling properties in cardiomyocytes treated with VAT EVs. **a,** Fluorescence signal representing change in membrane voltage of iPSC-aCMs during pacing at 1 Hz. Treatment of iPSC-aCMs with VAT-EVs (red) prolongs APD compared to control (black). **b,** IPSC-aCMs treated with VAT EVs (red, n = 21 wells from 4 VAT donors) have significantly prolonged APD compared to control (black, n = 15 wells, treated with iPSC-aCM-derived EVs). APD was measured during pacing at 1 Hz. **c,** Representative membrane voltage recordings of iPSC-vCMs, paced at 1 Hz, showing APD prolongation upon treatment with VAT-EVs (red) compared to control (black). **d,** IPSC-vCMs treated with VAT EVs (red, n = 19 wells from 5 VAT donors) have significantly prolonged APD compared to control (black, n = 7 wells, treated with iPSC-vCM-derived EVs). APD was measured during pacing at 1 Hz. **e,** Example of arrhythmia observed in paced iPSC-aCMs treated with VAT-EVs. **f,** IPSC-aCMs treated with VAT-EVs (n = 4 events out of 21 wells) have higher incidence of arrhythmia events (3 oscillations of voltage and 1 early afterdepolarization) compared to control (n = 0 events out of 15 wells). **g,** Fluorescence signal representing change in intracellular Ca^2+^ of iPSC-aCMs, paced at 0.5 Hz. Treatment of iPSC-aCMs with VAT EVs alters dynamics of Ca^2+^ transient. **h,** Ca^2+^ transient magnitude, transient time to peak (time to peak) and decay time (τ) in iPSC-aCMs treated with VAT EVs or control EVs. (red, n = 14 wells from 3 VAT donors), compared to control (black, n = 10 wells, treated with iPSC-aCM-derived EVs). **i,** Similar alteration of Ca^2+^ transient dynamics is observed in iPSC-vCMs treated with VAT EVs. **j,** Ca^2+^ transient magnitude, transient time to peak (time to peak) and decay time (τ) in iPSC-vCMs treated with VAT EVs or control EVs. (red, n = 16 wells from 3 VAT donors), compared to control (black, n = 10 wells, treated with iPSC-vCM-derived EVs). *, p < 0.05, **, p < 0.01, ***, p < 0.001.

Alteration of action potential shape and/or duration is a known risk factor for arrhythmias. In fact, we observed (pro)arrhythmic behaviors in iPSC-aCMs treated with VAT EVs, including early after depolarization and oscillation of membrane voltage independent of pacing stimulus (**Fig. 2e**). These (pro)arrhythmic events were observed more frequently in cardiomyocytes treated with VAT EVs compared to control (**Fig. 2f**) which likely corresponds to the observed increased risk of arrhythmia in obese individuals.

In addition to membrane voltage disturbances, dysfunction of intracellular Ca^2+^ cycling may also drive arrhythmogenesis. Therefore, we examined the effect of VAT EV treatment on intracellular Ca^2+^ transients in cardiomyocytes. **Fig. 2g** shows representative Ca^2+^ transient of iPSC-aCMs at 0.5 Hz pacing rate. IPSC-aCMs treated with VAT EVs demonstrate slower upstroke (time to peak) and slower decay time (τ) compared to control (**Fig. 2h**, time to peak 410±51 ms vs 342±23 ms, p = 0.0008; τ 362±41 ms vs 292±33 ms, p = 0.0001) although without significant difference in Ca^2+^ transient magnitude (normalized fluorescence intensity 0.92±0.17 vs 0.86±0.14, p = 0.37). Similar alteration in Ca^2+^ transient dynamics is observed in iPSC-vCMs treated with VAT EVs (**Fig. 2i-j**) where VAT EV treatment led to prolongation of upstroke (565±41 ms vs 499±26 ms, p = 0.02), slower decay time (403±87 ms vs 279±47 ms, p = 0.001), and no significant change in Ca^2+^ transient magnitude (0.70±0.14 vs 0.71±0.17, p = 0.77).

In addition to systolic Ca^2+^ level, diastolic Ca^2+^ and the sarcoplasmic reticulum (SR) Ca^2+^ content could also affect proarrhythmic behavior of cellular substrate. Therefore, we measured diastolic Ca^2+^ level and SR content in cardiomyocytes in treatment and control groups. Although there was impaired systolic Ca^2+^ cycling, there was no statistically significant difference in diastolic Ca^2+^ level or SR Ca^2+^ content in iPSC-aCMs and iPSC-vCMs treated with VAT EVs, compared to control (**Supplementary Fig. 4**). Disruption in overall Ca^2+^ transient dynamic can impact the well-orchestrated interaction between membrane voltage and intracellular Ca^2+^ during action potential generation leading to an increased risk of arrhythmogenesis.

### VAT EVs induce a pro-fibrotic and inflammatory state in non-cardiomyocyte cell types central to arrhythmogenesis *in vitro*

Apart from cardiomyocytes, cardiac-resident fibroblasts and macrophages are critical in the pathogenesis of arrhythmia through effects on inflammation and fibrosis. We first investigated whether VAT EVs influence fibroblast trans-differentiation into myofibroblasts—cells characterized by enhanced contractility and increased extracellular matrix production^27^ and a characteristic response of fibroblasts to injury or activation of the transforming growth factor beta 1 (TGF-β1) signaling pathway. Real-time quantitative PCR analysis revealed a significant upregulation of myofibroblast markers after treatment of fibroblasts with VAT EVs, including α-smooth muscle actin (*α-SMA*, 2.3±0.7 fold, p = 0.003) and periostin (7.3±3.8 fold, p = 0.005) in VAT EV-treated cells compared to controls (fibroblasts treated with the same concentration of fibroblast-derived EVs) and upregulation of collagen type 1 (*COL1A1*, 2.3±0.8 fold, p = 0.002), a major component of extracellular matrix (**Fig. 3a**). Additionally, iPSC-CFs treated with VAT EVs also secreted more collagen into culture media compared to control (**Fig. 3b**, 1.7±0.4 fold, p = 0.02). Overall, treatment with VAT EVs increased trans-differentiation of iPSC-CFs to myofibroblasts and increased collagen production, both of which are hallmarks of increased cardiac tissue fibrosis, a major risk factor for arrhythmogenesis.

**Figure 3:**
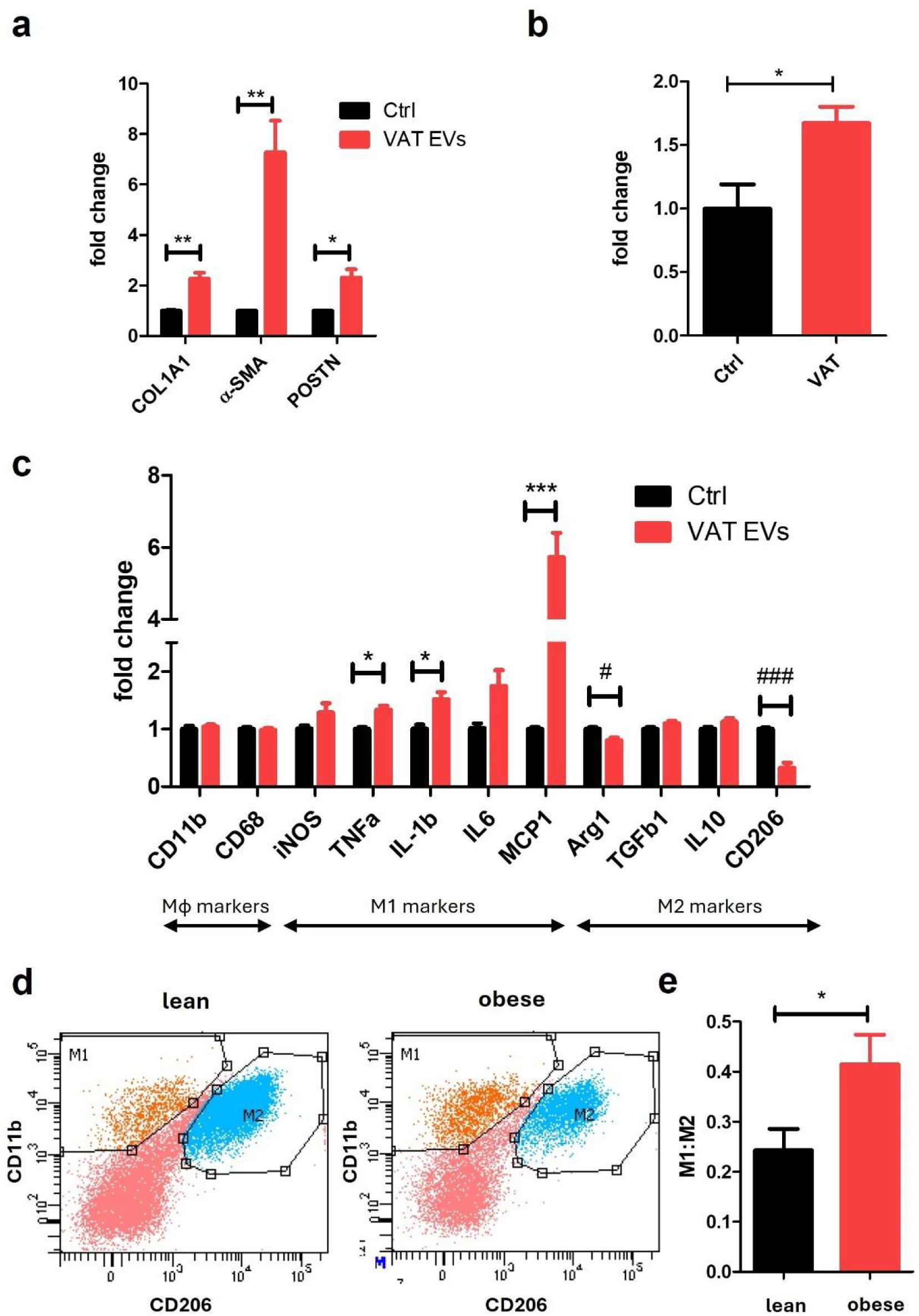
VAT EVs are pro-fibrotic and pro-inflammatory. **a,** Quantitative real-time PCR of iPSC-CFs demonstrates significant upregulation of markers of myofibroblasts upon treatment with VAT EVs (red, n = 11 wells from 4 VAT donors) compared to control (black, n = 5 wells, treated with iPSC-CF-derived EVs). *, p < 0.05/3, ** p < 0.01/3. **b,** IPSC-CFs treated with VAT-EVs secret significantly more collagen into culture media (red, n = 12 wells from 6 VAT donors) compared to control (black, n = 5 wells, treated with iPSC-CF-derived EVs). *, p < 0.05. **c,** Quantitative real-time PCR analysis of macrophage polarization markers in THP-1-derived macrophages treated with VAT EVs (red bars, n = 8 wells from 3 VAT donors) compared to control macrophages treated with macrophage-derived EVs (black bars, n = 5 wells). VAT EV treatment significantly upregulated M1 pro-inflammatory markers (TNF-α, IL-1β, IL-6, MCP-1) and downregulated M2 anti-inflammatory markers (Arg1, CD206) compared to control. Data presented as fold change relative to control ± SEM. *, p < 0.05/5, ***, p < 0.001/5 for M1 marker comparisons; #, p < 0.05/4, ###, p < 0.001/4 for M2 marker comparisons. **d,** Representative flow cytometry scatter plots showing macrophage populations based on CD11b and CD206 expression. Left panel shows macrophages from left ventricles of lean mice, right panel shows macrophages from left ventricles of obese mice. **e,** Quantification of M1:M2 ratio from flow cytometry analysis comparing lean (black bar) and obese (red bar) conditions. Compared to lean, obese mice have significant increase in M1:M2 ratio, indicating enhanced pro-inflammatory polarization. Data presented as mean ± SEM. *p < 0.05.

Next, we investigated the effect of VAT EVs on macrophage polarization toward pro-inflammatory phenotypes. **Fig. 3c** shows increased expression of markers associated with a pro-inflammatory M1 phenotype, including *TNF-α* (1.3±0.2 fold, p = 0.004), *IL-1β* (1.5±0.3 fold, p = 0.008), and *MCP-1* (5.7±1.8 fold, p = 0.0002), and decreased expression of anti-inflammatory M2 phenotype, including *Arg1* (0.8±0.1 fold, p = 0.007) and *CD206* (0.3±0.2 fold, p = 0.0001), upon VAT EV treatment. Additionally, flow cytometry analysis of cardiac macrophages in lean versus obese mouse hearts confirmed these findings, showing increased M1:M2 ratios in obese compared to lean conditions (**Fig. 3d-e**, 0.4±0.2 vs 0.2±0.4, p = 0.038). This pro-inflammatory response of cardiac macrophages could create a vulnerable cardiac substrate conducive to arrhythmogenesis.

### *In vivo* evidence of adipose to myocardial communication by adipose-derived EVs

Given the *in vitro* effects of VAT EVs on cardiomyocytes, we next studied whether adipose-derived EVs reach the heart *in vivo* in obesity. We employed a unique transgenic mouse model, *Exomap1*, where tissue-specific EVs are tagged with green fluorescent protein and human CD81 surface marker protein in response to Cre expression^26^, which allows tracking of their biodistribution. Crossing this Exomap1 transgenic line with Adiopoq-Cre transgenic mice, where a Cre recombinase is expressed under control of adiponectin promoter/enhancer region, allows tracking of tagged adipose tissue specific green EVs. Only double positive Adipoq-Cre^+^/Exomap^+^ mice expressed a fusion protein hsCD81-mNeonGreen, as evident by green fluorescence signal from inguinal white adipose tissue (iWAT) and immunoblot of iWAT lysate against mNeonGreen protein (**Fig. 4a-c** and **Fig. 5a**).

**Figure 4:**
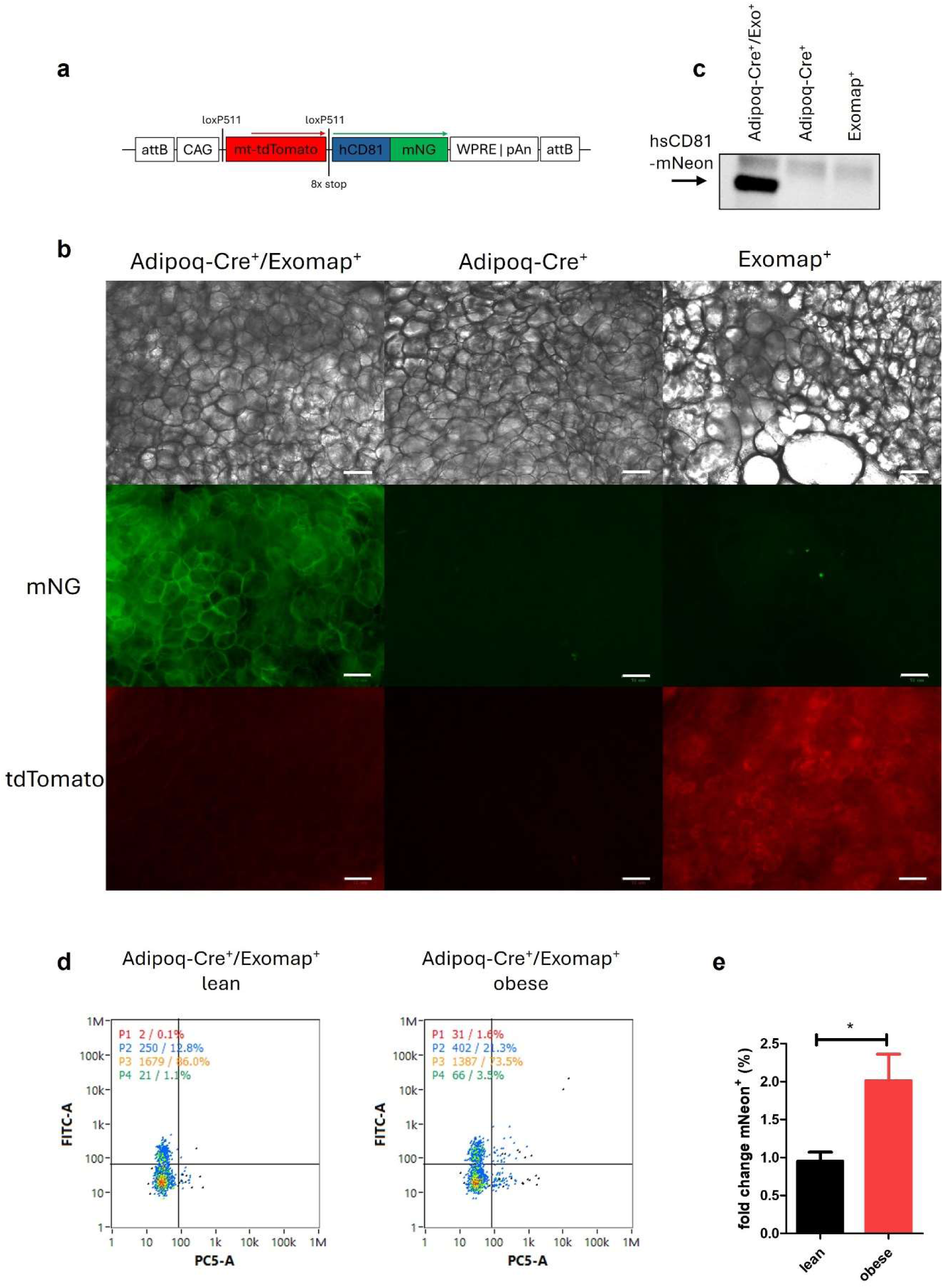
Characterization of Adipoq-Cre^+^/Exomap^+^ mice. **a,** Line diagram displaying transfer vector (top) used to create *Exomap* mouse line (adapted from Fordjour et al^26^). The Exomap mouse line was crossed with *Adipoq-Cre* line (Adiopoq) where a Cre recombinase is expressed under control of adiponectin promoter/enhancer region. In the absence of Cre recombinase, mitochondrial red fluorescence protein (mt-tdTomato) is expressed. In the presence of Cre recombinase, green fluorescent protein (mNeoGreen)-tagged human CD81 (hCD81-mNG) is expressed and localized to the plasma membrane. **b,** Bright field (top), and fluorescence images (middle, green channel, bottom, red channel) of freshly mounted inguinal white adipose tissue (iWAT). Only iWAT from double positive mouse (Adipoq-Cre^+^/Exomap^+^, left column), express hCD81-mNG on plasma membrane with absence of red mt-tdTomato fluorescence, while IWAT of Adipoq-Cre^+^ line (middle column) does not contain either green hCD81-mNG or red mt-tdTomato fluorescence signal. IWAT of Exomap^+^ mouse only expresses red mt-tdTomato fluorescence. Scale bar represents 72 μm. **c,** Western blot of iWAT lysates from Adipoq-Cre^+^/Exomap^+^, Adipoq-Cre^+^, and Exomap^+^ mouse probed with anti-mNeonGreen antibody. **d,** Nano Flow Cytometry analysis of circulating sEVs of lean and obese Adipoq-Cre^+^/Exomap^+^ mice. Circulating sEVs of Adipoq-Cre^+^/Exomap^+^ mice contain a surface marker hCD81, tagged with green mNeonGreen fluorescence protein. Plasma of both lean and obese Adipoq-Cre^+^/Exomap^+^ mice contain adipose tissue specific hCD81-mNG-tagged sEVs. **e,** Plasma of obese Adipoq-Cre^+^/Exomap^+^ mice contain higher number of adipose tissue specific hCD81-mNG-tagged sEVs compared to lean mice (n = 4 lean, 3 obese, p < 0.05).

**Figure 5:**
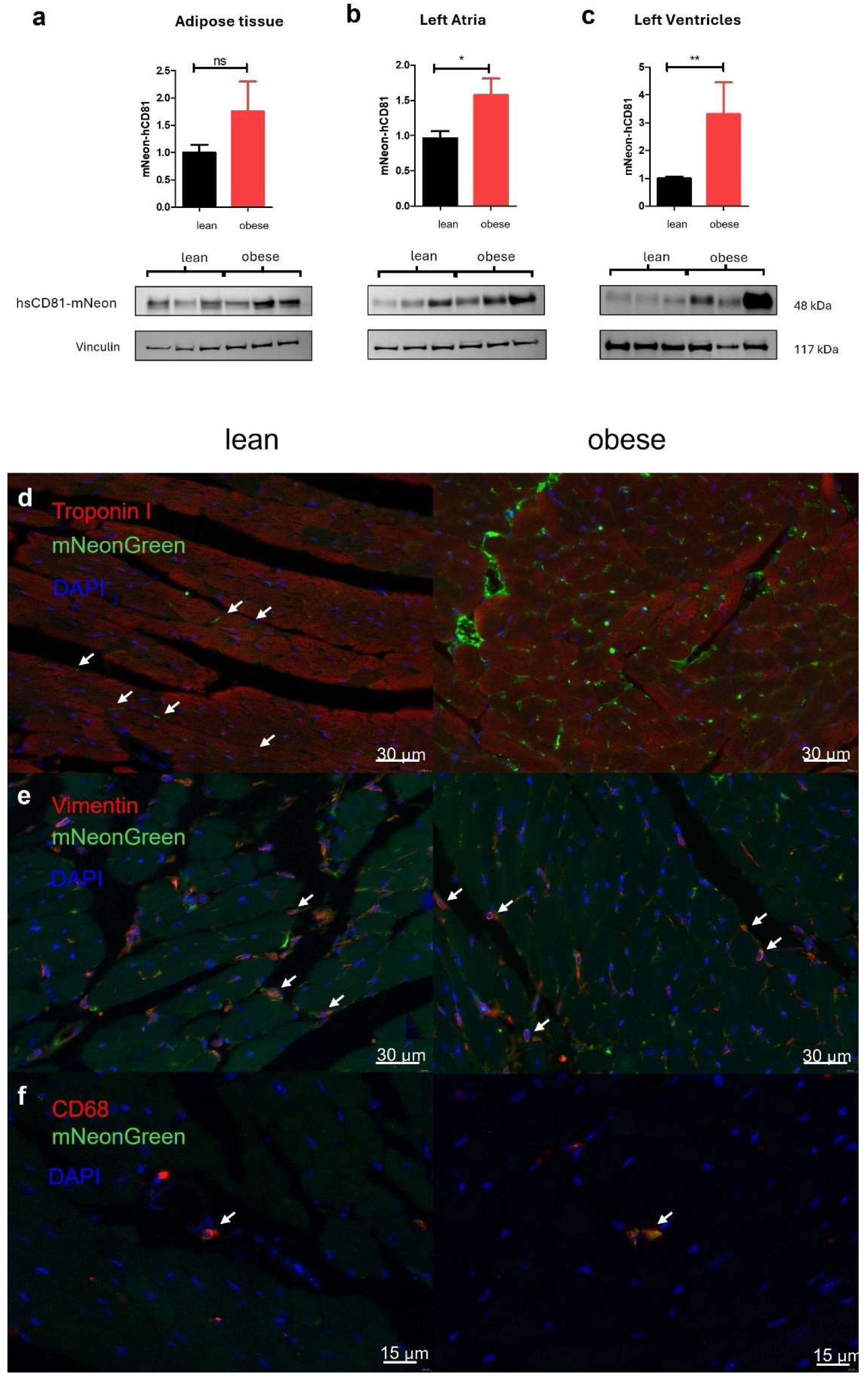
Adipose tissue-derived EVs are deposited in the hearts. **a,** Western blot of iWAT lysate probed with anti-mNeonGreen antibody demonstrates presence of hCD81-mNeonGreen protein in iWAT tissue of both lean and obese mice (bottom row), indicating successful Cre recombination and expression of hCD81-mNeonGreen recombinant protein in iWAT tissue. There is no significant difference between normalized amount of hCD81-mNeonGreen (using vinculin as a loading control) in iWAT between lean and obese groups. **b,** Western blot of lysate from left atria probed with anti-mNeonGreen antibody demonstrates presence of hCD81-mNeonGreen of both lean and obese mice (bottom row), indicating presence of adipose tissue-specific EVs tagged with hCD81-mNeonGreen. **c,** Western blot of lysate from left ventricles probed with anti-mNeonGreen antibody (bottom row) demonstrates presence of adipose tissue-specific EVs tagged with hCD81-mNeonGreen in left ventricle (bottom row). Normalized amount of hCD81-mNeonGreen is significantly higher in left ventricles of obese compared to lean mice. (n = 3 lean mice and 3 obese mice, each with 2 technical replicates) *, p < 0.05, **, p < 0.01. **d-f,** Immunostaining of left ventricles of lean and obese mice demonstrated co-localization of troponin I, a cardiomyocyte marker (**d**), vimentin, a fibroblast marker (**e**), CD68, a macrophage marker (**f**), and mNeonGreen, indicating adipose tissue-specific EV deposition in these three cell types.

After 16 weeks, the body weight of double positive Adipoq-Cre^+^/Exomap^+^ male mice fed with high fat diet (“obese”) was significantly higher than those fed with regular diet (“lean,” **Supplementary Fig. 5a**). However, there was no significant difference in the weight of the ventricles normalized by tibial length (**Supplementary Fig. 5b**) or the degree of histological cardiac fibrosis (**Supplementary Fig. 5c-d**) between the two groups. To establish communication between adipose and cardiac tissue, we first demonstrated that adipose tissue-specific EVs, tagged with hsCD81-mNeonGreen protein, are secreted into plasma (**Fig. 4d**) and consistent with many prior studies^21,28^, obesity was associated with a higher proportion of adipose-derived circulating EVs (**Fig. 4e**). Next, to establish that circulating adipose-derived EVs target the heart, we performed immunoblot of cardiac tissue lysate which demonstrated presence of hsCD81-mNeonGreen protein in both left atria (LA) and left ventricles (LV), indicating that adipose tissue-specific EVs were indeed trafficked to the heart (**Fig. 5b-c**). Interestingly, both LA and LV from obese mice had significantly higher amount of deposited adipose tissue-specific EVs compared to lean mice, as evident by higher intensity of normalized hsCD81-mNeonGreen bands to a housekeeping protein, vinculin, in immunoblot (1.6±0.6 fold, p = 0.039 for LA and 3.43±2.8 fold, p = 0.004 for LV).

Additionally, immunohistochemistry of left ventricular tissue showed that adipose tissue-specific EVs are deposited in cardiomyocytes, cardiac fibroblasts, and macrophages, as evident by double positive staining of mNeonGreen and cardiac troponin I, vimentin, and CD68, respectively (**Fig. 5d-f**). In vivo demonstration that the three pillars of arrhythmogenic substate are targets of adipose-derived EVs validates the physiological relevance of our in vitro findings.

### VAT EVs induce broad alterations in transcriptional profile of cardiomyocytes, fibroblasts, and macrophages *in vitro*

To further explore potential mechanisms of VAT-EV-associated arrhythmogenesis, we next performed RNA sequencing of VAT-EV-treated cardiomyocytes, fibroblasts, and macrophages (**Fig. 6a-h**). Genes involved in electrical and structural remodeling, inflammation, and cholesterol and lipid metabolism were prominently affected in both atrial and ventricular cardiomyocytes (**Supplementary Fig. 6**). In atrial cardiomyocytes (**Fig. 6b** and **Supplementary Table 4**), we observed alteration in gene expression of ion channels (*CACNA1G, HCN4, SCN9A, TRPC3*), calcium handling proteins (*RYR3, TRPC3*), structural elements (*CNTN5, ACTA2*, various collagen types), natriuretic peptides indicative of hemodynamic stress (*NPPA, NPPB*), and cholesterol/lipid metabolism (*HMGCR, LDLR*). Ventricular cardiomyocytes (**Fig. 6d** and **Supplementary Table 5**) displayed upregulation of genes involved in excitation-contraction coupling and metabolic remodeling (93 overlapping differentially expressed genes with atrial cardiomyocytes, **Supplementary Table 6**).

**Figure 6:**
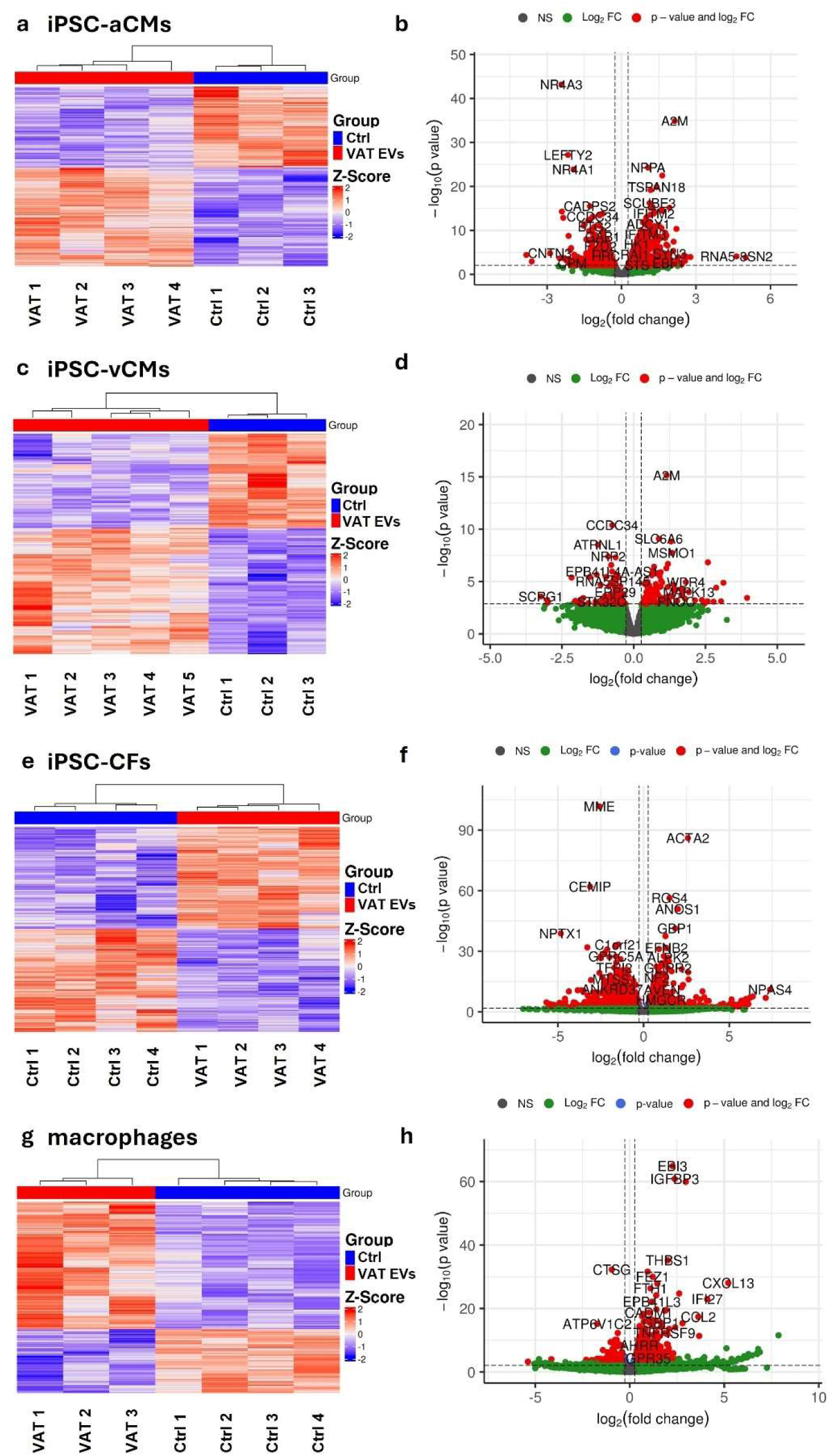
Treatment with VAT EVs alter transcriptional landscape. Bulk RNA sequencing of iPSC-aCMs (**a-b**), iPSC-vCMs (**c-d**), iPSC-CFs (**e-f**), and macrophages (**g-h**), comparing VAT EV-treated groups to their respective controls. **(a, c, e, g)** Hierarchical clustering was performed on control (Ctrl) and treatment (VAT EVs) groups based on the differentially expressed genes. The horizontal axis is composed of all the samples in both groups, and vertical axis includes all differentially expressed genes. Top, control samples are denoted in blue squares and treatment samples in red squares. Dark blue to dark red color gradient illustrates lower to higher expression. **(b, d, f, h)** Volcano plot was created by all differentially expressed genes. The y axis shows the FDR-adjusted p value, and the x axis displays the log2 fold-change value. The red dots represent the differentially expressed genes with FDR-adjusted p ≤ 0.1 and absolute fold-change ≥ 1.2, while green dots represent non-significantly differentially expressed genes.

VAT-EV treatment induced significant changes in genes in cardiac fibroblasts related to extracellular matrix production and myofibroblast differentiation (**Fig. 6f** and **Supplementary Table. 7**). Collagen genes (*COL1A1, COL3A1*) were strongly affected (*COL1A1*, 8-fold upregulation), alongside myofibroblast markers (*ACTA2, POSTN*). Macrophages showed strong upregulation of inflammatory pathways and downregulation of anti-inflammatory mediators, consistent with our functional observations of M1 polarization (**Fig. 6h** and **Supplementary Table 8**).

### Transcriptome-wide genetic association studies implicate genes induced by VAT-EVs in arrhythmogenesis

We next employed transcriptome-wide association studies (TWAS) in left ventricle and atrial appendage to perform genetic inference on key arrhythmic phenotypes in the heart (QT interval and AF) for the genes dysregulated by VAT-EVs. TWAS revealed significant association (p < 5.56 x 10^-5^) of 32 unique differentially expressed genes with QT interval and 24 associated with AF (**Supplementary Table 9**). The QT interval TWAS (**Fig. 7a**) identified several genes with established roles in cardiac repolarization that were significantly altered by VAT-EV treatment. For example, *STRN*, associated with cardiac hypertrophy and structural remodeling, showed significant upregulation in both atrial and ventricular cardiomyocytes. *TMEM176*, which plays a role in inflammasome activation and has been linked to arrhythmia susceptibility, was prominently upregulated across both cell types. *FADS1*, involved in lipid metabolism and known to influence membrane composition and ion channel function, showed differential expression patterns. *SYNPO2L*, important for sarcomere development and contractile function, was also significantly affected.

**Figure 7:**
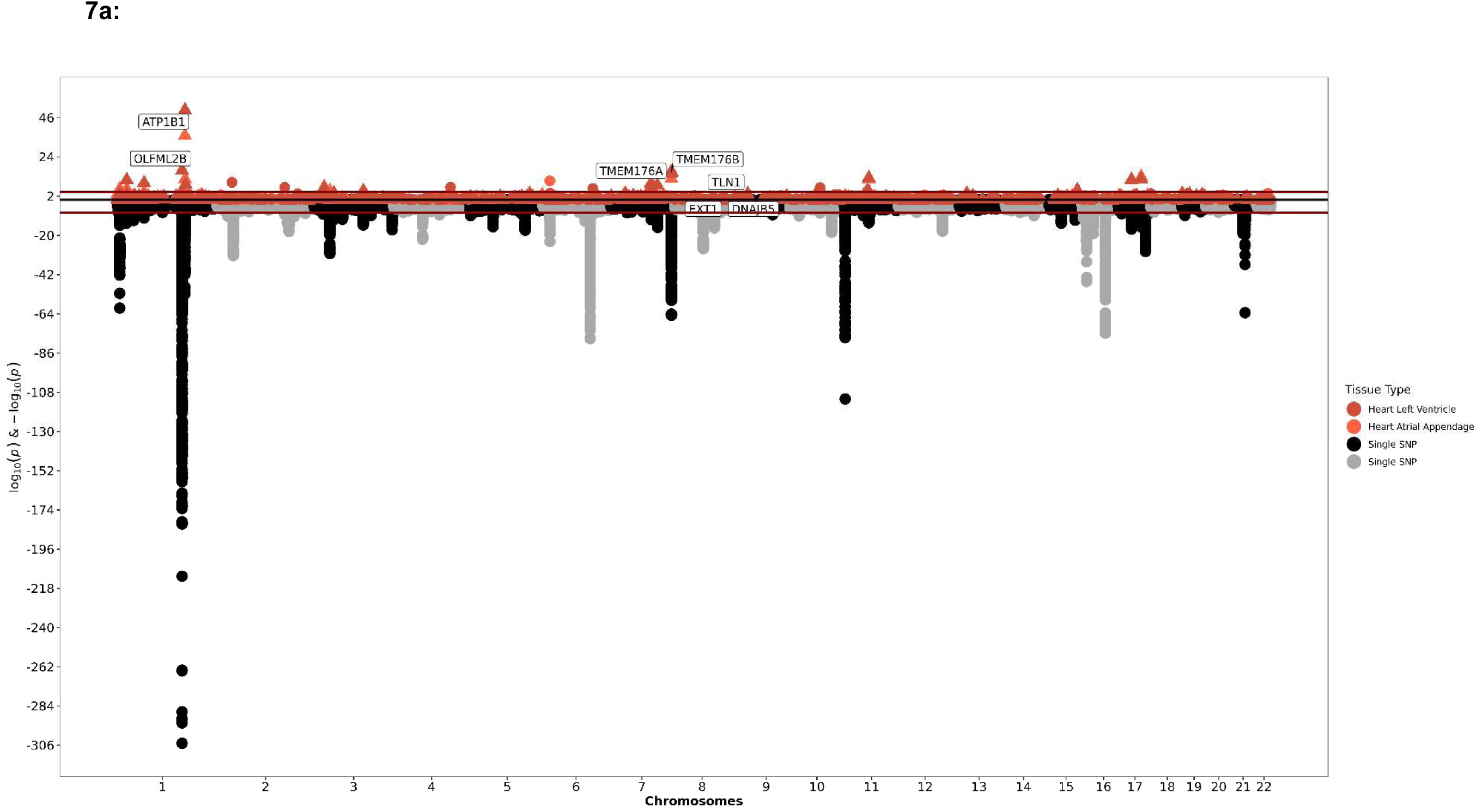
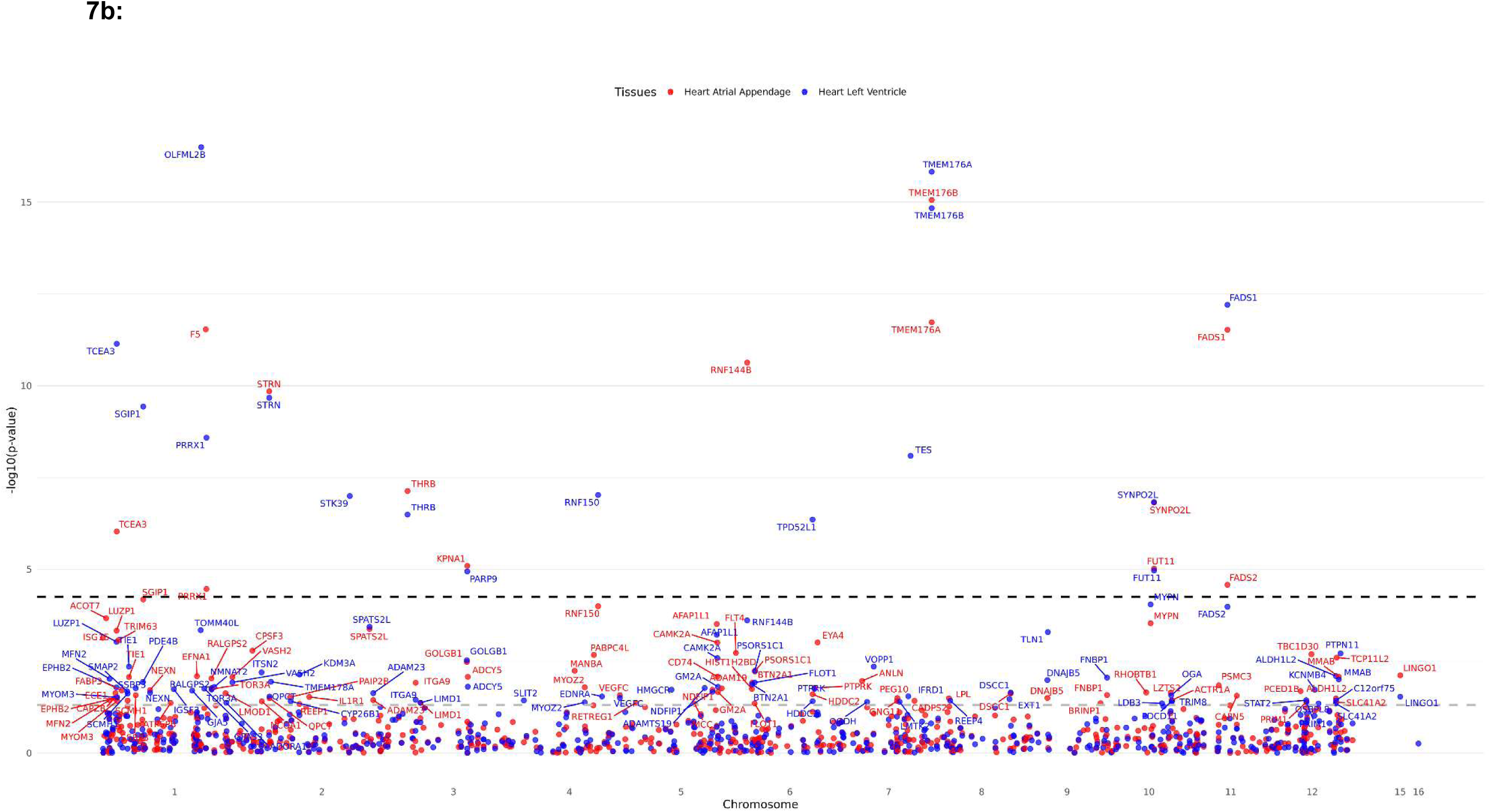
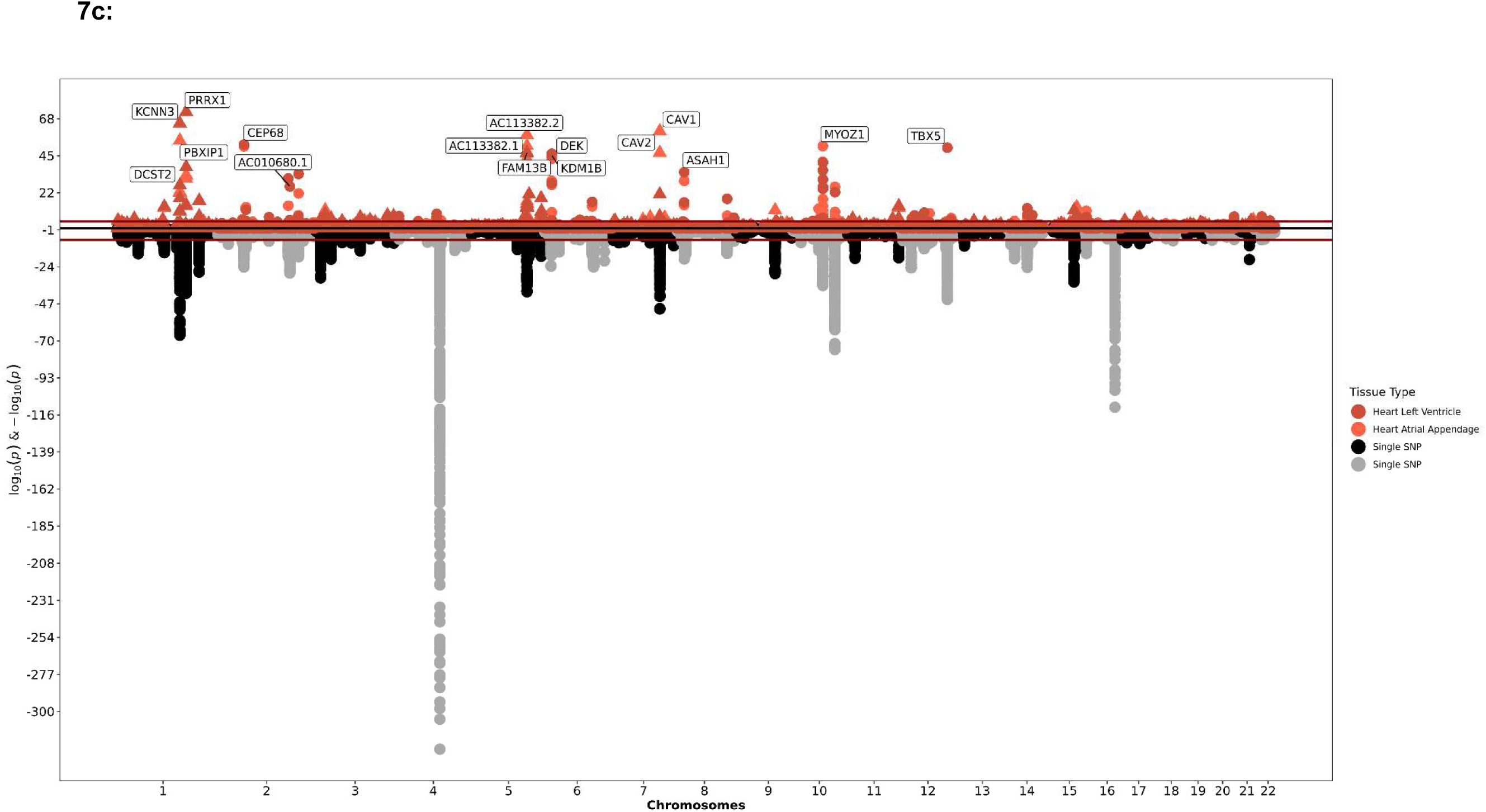
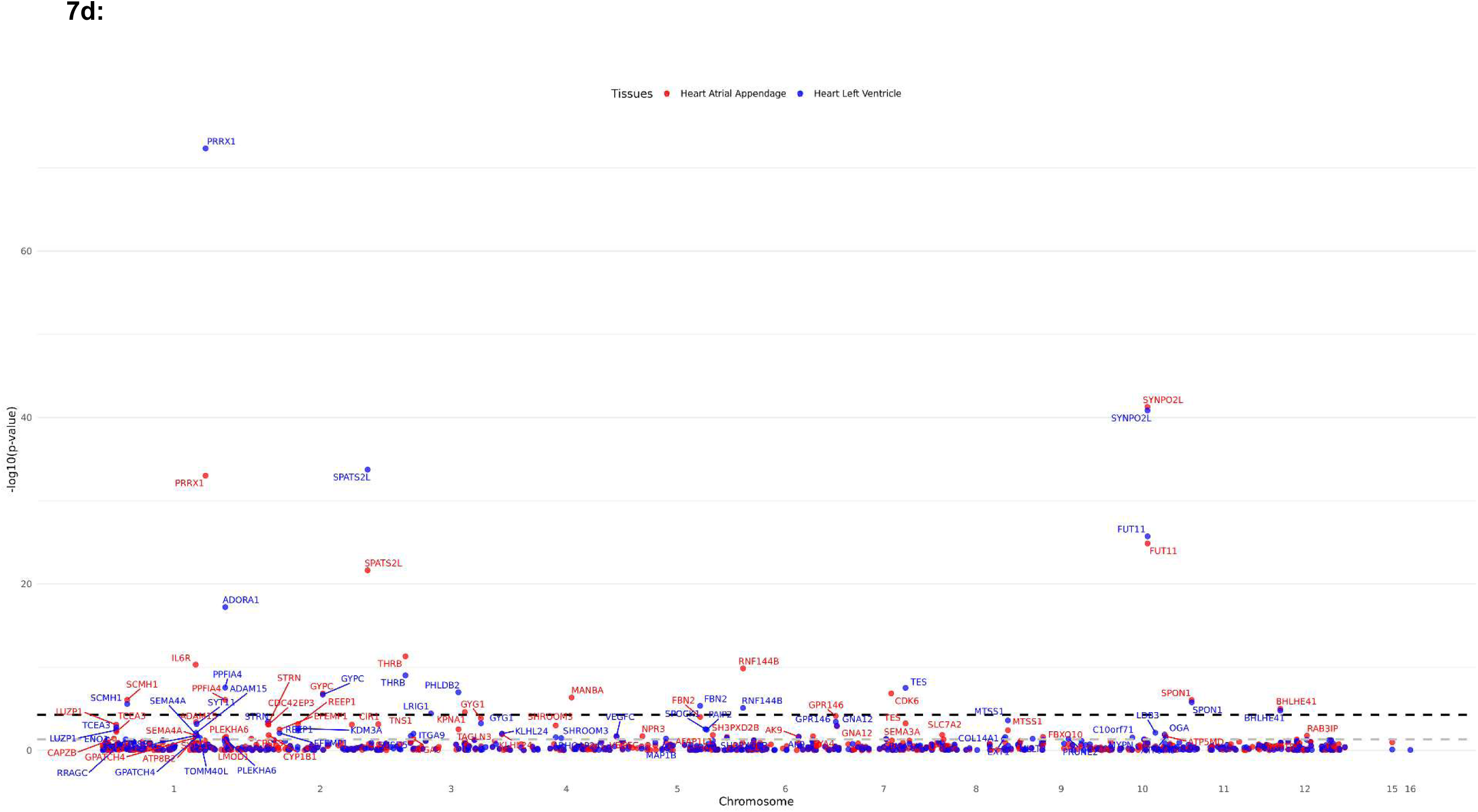
TWAS of QT interval and atrial fibrillation. **a,** TWAS of QT interval identified 32 genes that attained genome-wide significance. The red line above indicates genome-wide TWAS significance (p < 5.56 x 10^-5^), and the red line below indicates genome-wide GWAS significance (p < 5 x 10^-8^). **b,** Eleven unique QT interval-associated genes were differentially expressed in *in vitro* studies and linked (via Hi-C) to a genome-wide significant (p < 5 x 10^-8^) regulatory GWAS SNP in the same cell type or tissue in which differential expression was observed (atrium, ventricle, fibroblast, or macrophage). The threshold for genome-wide significance (*p* < 5 x 10^-8^) is denoted in black, and the threshold for nominal significance (p < 0.05) is denoted in gray. **c,** TWAS of atrial fibrillation identified 24 genes that attained genome-wide significance. **d,** Atrial fibrillation TWAS associations with differentially expressed genes in *in vitro* studies and linked (via Hi-C) to a genome-wide significant (p < 5 x 10^-8^) regulatory GWAS SNP in the same cell type or tissue in which differential expression was observed (atrium, ventricle, fibroblast, or macrophage). We identified 20 unique genome-wide significant (p < 5.56 x 10^-5^) genes that met these criteria.

Since TWAS was performed in atrial and ventricular bulk tissue, inference on whether specific genes in a specific cell type (e.g., atrial and ventricular cardiomyocyte, fibroblasts, macrophages) are related to our phenotype is inaccessible to standard TWAS methodology. Therefore, we utilized cell-type-specific epigenomic, chromatin accessibility, and chromatin conformation assays to provide <u>cell-type-specific</u> genetic evidence linking QT interval with the VAT-EV induced gene expression changes in the four cell types tested. We identified a subset of 11 genes in contact (via Hi-C) with a *cis*-regulatory element in the same cell type in which differential expression was observed (atrial cardiomyocyte, ventricular cardiomyocyte, fibroblast, or macrophage); this *cis*-regulatory element overlapped with a SNP associated with QT interval at genome-wide significance (**Fig. 7b** and **Supplementary Table 10**).

We found similar results in our AF TWAS (**Fig. 7c**), including *PRRX1* (directly linked to familial AF in humans^29^), *LRRC10* (implicated in dilated cardiomyopathy and alter gating of L-type Ca2+ current^30^), *ADORA1* (modulate heart rate and conduction velocity through modulations of multiple ion channels^31,32^), and *SYNPO2L* (sarcomere development^33^). Analogous to our epigenetic and chromatin-based studies with QT interval, of the 24 differentially expressed genes showing significant association with AF in TWAS, 20 were in contact (via Hi-C) with a *cis*-regulatory element in the same cell type where we observed differential expression (atrial cardiomyocyte, ventricular cardiomyocyte, fibroblast, or macrophage); again, this *cis*-regulatory element overlapped with a SNP associated with AF at genome-wide significance (**Fig. 7d**). These results provide genetic support linking the effects of VAT-EVs on key cell types in the heart to key obesity-driven arrhythmic risk.

### *TRPC3* mediates VAT-EV-induced action potential prolongation

As a proof-of-principle for the potential for adipose-heart communication via EVs to identify modifiable targets in arrhythmias, we focused on modulating *TRPC3* (a cardiac ion channel upregulated after VAT-EV exposure in cardiomyocytes). TRPC3 channels conduct non-selective cation currents that are upregulated during stress and can prolong APD when overexpressed^34^ but have not previously been known to be involved in the pathogenesis of AF. TRPC3 was significantly upregulated in atrial myocytes (2.7-fold, FDR-adjusted p-value 2.8×10^-6^) upon VAT EV treatment. We co-treated cardiomyocytes with VAT-EVs and Pyr3, a small molecule selective TRPC3 inhibitor^35^. In atrial cardiomyocytes (**Fig. 8a-b**), Pyr3 treatment abrogated VAT-EV-induced APD prolongation, with APD_80_ values returning to <u>near-baseline</u> (301 ± 12 ms with Pyr3 vs 396 ± 8 ms without Pyr3, vs 345 ± 6 ms in untreated controls, p<0.0001). In ventricular cardiomyocytes (**Fig. 8c-d**), we observed a similar rescue, with Pyr3 treatment reducing APD_80_ from 602 ± 13 ms to 540 ± 7 ms (p = 0.02).

**Figure 8:**
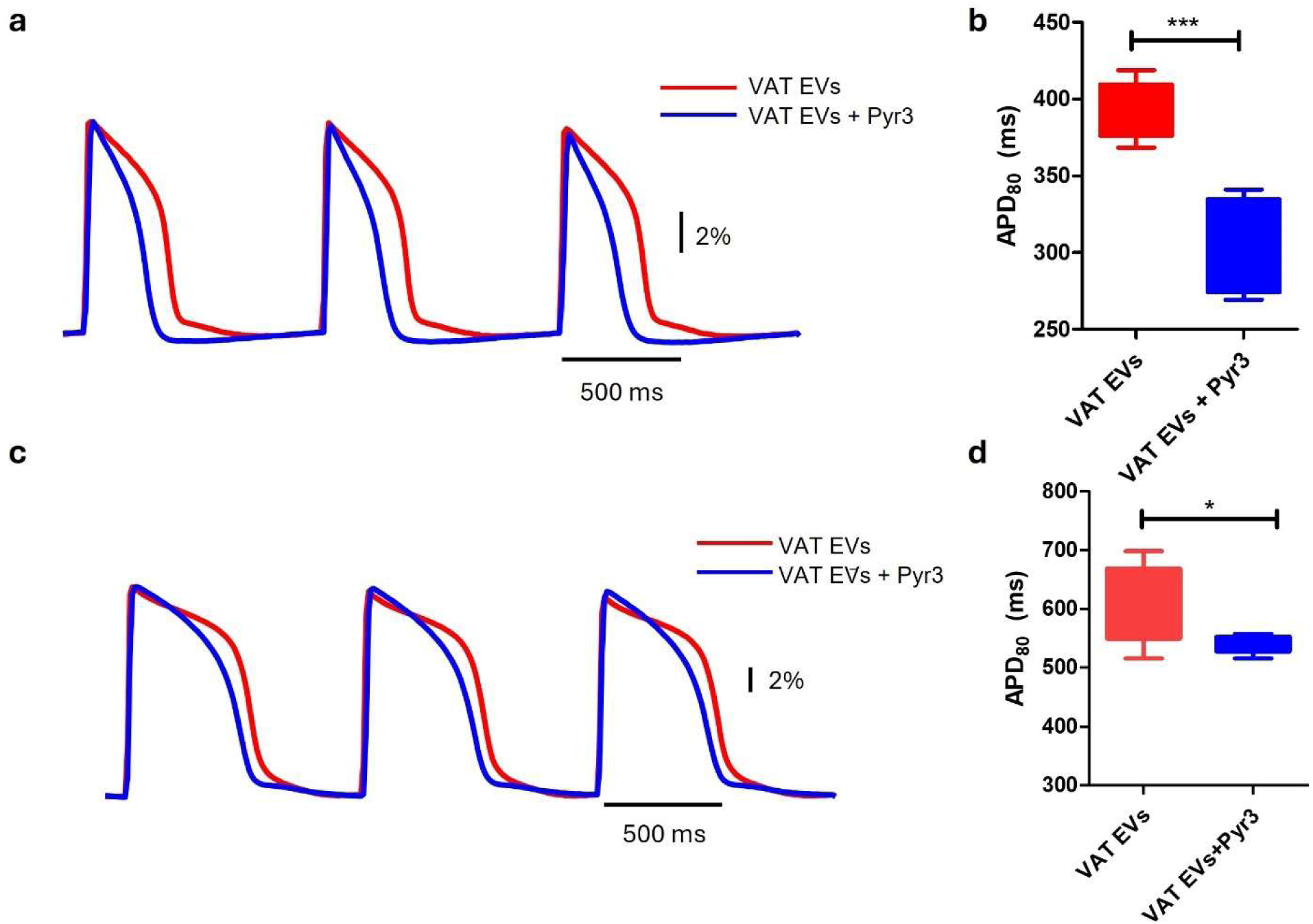
TRPC3 inhibition rescues VAT EV-induced APD prolongation. **a,** Representative action potential traces and quantification in atrial cardiomyocytes showing complete rescue of APD prolongation by Pyr3 treatment. **b**, The action potential morphology is restored to near-control levels, with normalized repolarization kinetics. (red, VAT EVs n = 6 wells; blue, VAT EVs + Pyr3, n = 6 wells) **c-d,** Similar analysis in ventricular cardiomyocytes demonstrating significant rescue of APD prolongation by TRPC3 inhibition (red, VAT EVs n = 19 wells; blue, VAT EVs + Pyr3, n = 5 wells) *, p < 0.05, ***, p < 0.001.

## DISCUSSION

The last several decades have observed a parallel increase in the prevalence of AF and obesity^36^. Traditionally, obesity increases cardiovascular risk through effects on systemic inflammation and metabolic stress^37^, though recent work suggests that local adipose mediators may play a role in key obesity-related cardiac complications^20,38^. Whether and how adipose tissue “signals” to the heart in obesity remains open. Here, we present evidence linking VAT EVs from obese individuals to cardiac arrhythmia pathogenesis via integrated model system, cellular, and genetic approaches. We found that treatment of iPSC-CMs with VAT EVs phenocopies APD prolongation observed in human atrial tissue from obese individuals and produces broad shifts in the transcriptional landscape of human iPSC-derived atrial and ventricular cardiomyocytes. These differentially expressed genes displayed causal association with AF and ventricular arrhythmia susceptibility via transcriptome-wide genetic association methods in large GWASs of atrial fibrillation and QT interval. Furthermore, the effects of VAT EVs extended to pro-inflammatory macrophage polarization and pro-fibrotic fibroblast activation, key steps in lowering arrhythmic thresholds in the heart. Importantly, we observed adipose-myocardial crosstalk *in vivo* through genetically engineered mouse models of EV tracking, specifically in obese rodents. Finally, focusing on *TRPC3* (a pharmacologically targetable ion channel gene differentially expressed on VAT EV exposure), we demonstrated abrogation of the effect of the VAT EVs on APD with a selective small molecule TRPC3 inhibitor, providing additional evidence linking adipose-heart EV communication and obesity-induced AF. Collectively, through comprehensive molecular genetics, human, and experimental evidence, these results underscore the importance of direct adipose-myocardial communication as a targetable mechanism of arrhythmogenesis in obesity.

Individuals with obesity have a heightened risk for both atrial and ventricular arrhythmia, thought to be mediated by a panoply of systemic inflammatory factors^19,39^, adverse cardiac remodeling^40,41^, and cardiometabolic comorbidity^42,43^. Recent data across over 50,000 individuals worldwide suggests shared genetic links between obesity and AF^16^, though how obesity <u>itself</u> (not only via its comorbidities) impacts arrhythmia is less clear. In this space, EVs have emerged as endocrine transducers of molecular signals across tissues in obesity-related complications. Adipose tissue serves an important source of circulating EVs^28^, with weight loss causing a reduction in the number of overall and adipose-derived EVs in circulation as well as a shift in their cargo^44–46^. In addition, adipose-derived EVs impact broad inflammatory states relevant to metabolic health^47–49^, including macrophage polarization, hepatic fibrosis, muscle metabolism, and cardiovascular states. Most data in human arrhythmogenesis arises from explanted tissue from patients with established disease, demonstrating greater EV release and more pro-inflammatory/fibrotic contents from adipose tissue in patients with AF (relative to without AF)^50^, with the ability of these EVs to transfer these arrhythmic effects across cells. Specifically, transcriptional cargo (miRNAs) from these EVs can impact the membrane potential and potassium channel expression in rodent cardiomyocytes^51^.

The current report adds substantially to these findings. First, in contrast to APD shortening observed in most animal systems of arrhythmia susceptibility used in discovery efforts and in recent epicardial fat-derived EV studies^50^, we *directly* observed APD <u>prolongation</u> in electrophysiologic studies from human right atrial tissue in individuals <u>with obesity</u> (relative to non-obese), a phenotype reproduced upon treatment of iPSC-CMs with VAT EVs from individuals with obesity. This mechanism represents a departure from traditional models of AF (typically focus on APD shortening) and therapeutics (focus on *causing* APD prolongation). In addition, the impaired calcium handling we observed—specifically the prolonged time-to-peak and decay kinetics—further supports arrhythmogenic potential of VAT EVs: while peak calcium amplitude remained unchanged, the altered kinetics suggest disrupted excitation-contraction coupling that could contribute to both contractile dysfunction and electrical instability. Our data is a key contribution to the growing literature on differences in obesity-associated AF compared to AF associated with other conditions^52,53^.

In addition, our results extend previous proteomic results in adipose-derived EVs^50^, demonstrating a broad adverse impact of VAT EVs on cardiomyocytes, macrophages, and fibroblasts—key cells involved in preparing myocardial substrate for arrhythmia. The profibrotic response we observed in cardiac fibroblasts—characterized by upregulation of myofibroblast markers and increased collagen secretion—creates a substrate conducive to arrhythmogenesis through several mechanisms. Increased tissue fibrosis leads to conduction heterogeneity, slowed propagation velocity, and enhanced susceptibility to reentrant arrhythmias. The inflammatory activation of macrophages toward an M1 phenotype further compounds this proarrhythmic environment by promoting tissue remodeling and potentially affecting cardiomyocyte function through paracrine signaling^25^.

Furthermore, the overlap of genes differentially expressed upon VAT EV treatment in these target cells with large-scale GWASs of atrial fibrillation and QT interval emphasized the causal role of adipose-derived EVs on arrhythmia pathogenesis. Our approach improves upon standard TWAS methodology by incorporating epigenetic and chromatin-based studies in the target cell types to enable cell-type-specific genetic inference linking the genes differentially expressed upon VAT EV exposure to arrhythmia susceptibility. Key genes identified include *STRN* (associated with cardiac hypertrophy), *TMEM176* (involved in inflammasome activation), *FADS1* (lipid metabolism), and *SYNPO2L* (sarcomere development). The enrichment of transcription factors like *PRRX1*, which regulates cardiac development and fibrosis, suggests that VAT EVs may induce long-term epigenetic changes in cardiac cells. Finally, we observed for the first time *in vivo* <u>direct</u> communication between adipose tissue and the myocardium using a novel genetically engineered tracking system that labeled only adipose-derived EVs. The collection of our data—evidence of trans-organ communication, transfer of phenotype, and change in molecular state with reversibility (with TRPC3 modification)—provides compelling evidence to expand the current framework of obesity-induced arrhythmia to include trans-organ EV communication.

The identification of TRPC3 upregulation in VAT EV-treated cardiomyocytes through RNA sequencing exemplifies how comprehensive transcriptomic analysis can guide rational drug repurposing strategies. The fact that TRPC3 emerged as a significantly differentially expressed gene provided the mechanistic insight that led us to test Pyr3, an existing TRPC3 inhibitor originally developed for other applications. This approach represents a paradigm shift from traditional empirical drug screening to hypothesis-driven therapeutic targeting based on pathway-specific perturbations. The successful rescue of VAT EV-induced APD prolongation by Pyr3 treatment validates this transcriptomics-to-therapeutics pipeline and highlights the potential for systematically mining EV-induced gene expression changes to prioritize existing drug libraries for repurposing. This strategy could be particularly valuable for complex diseases like obesity-associated cardiovascular complications, where multiple pathways are simultaneously perturbed.

Several limitations of this study should be acknowledged. While our primary studies were on iPSC-CM models that may not fully recapitulate *in vivo* human states, the concordance between human right atrial tissue electrophysiologic studies and iPSC-CMs in APD is reassuring. We cultured all cell types separately with VAT EVs, given the complexity inherent in studying multi-cellular interactions that are likely at play *in vivo*. Future studies using co-culture systems, “heart on a chip” technologies^54^, or other organoids will likely more closely recapitulate how adipose-derived EVs impact the multicellular environment of the human heart. Other areas of study within adipose-tissue EVs (proteomics, non-coding RNA) beyond transcripts are an additional future area of investigation. Finally, the temporal dynamics of EV-mediated cardiac remodeling require further investigation: our acute treatment studies provide a snapshot of EV effects, but chronic exposure patterns (especially reversibility of EV-induced changes) will need to be characterized to inform therapeutic strategies.

In conclusion, our findings support a <u>direct</u> communication pathway between adipose tissue and the heart that promotes proarrhythmic remodeling. Our results challenge the paradigm of APD shortening as a major consequence of obesity in humans. *In vivo* genetic tracking models support transfer of EVs from adipose tissue to myocardium, and cellular studies in iPSC models support broad transcriptional and functional effects of adipose-derived EVs on pathophenotypes relevant to arrhythmia. Genetic studies (large scale TWAS) and functional studies (TRPC3 modulation) support a causal role for VAT EVs on arrhythmia susceptibility. Collectively, this work offers a new paradigm for obesity-related cardiovascular consequences, extending beyond traditional hemodynamic and metabolic factors to direct intercellular communication mechanisms that actively reprogram cardiac tissue toward a proarrhythmic state. Understanding and targeting these pathways offers a new pathway for therapeutic development amidst a growing burden of obesity-associated arrhythmia in an increasingly obesogenic environment.

## METHODS

### Human myocardial tissue

All procedures adhered to the principles of the Declaration of Helsinki and received approval from the ethics committees of the University of Regensburg and the University of Göttingen (approval numbers 14/9/11 and 21/10/00). Written informed consent was obtained from every participant. Samples of human atrial myocardium were collected during atrial resections performed during open-heart surgery (see **Supplementary Table 1** for patient details). Following removal, the tissue was immediately transferred into cardioplegic solution cooled to 4 °C for transport.

### Human cardiomyocytes isolation

Atrial myocardium from patients was used for cellular experiments. Cardiomyocyte isolation was preformed based on an established method by Pabel et al^55^. Prior to isolation, atrial tissue was carefully cleaned of adipose tissue and blood vessels, minced into fine fragments, and rinsed thoroughly. The samples were washed three times in a Ca²⁺-free buffer composed of (mM): NaCl 100, glucose 20, KCl 10, KH₂PO₄ 1.2, MgCl₂ 5, taurine 50, BDM 10, and MOPS 5 (pH 7.2). Tissue fragments were then incubated at 37 °C in a spinner flask with continuous oxygenation, containing Ca²⁺-free buffer supplemented with 0.77 mg/mL collagenase (Worthington type 1, 370 U/mg) and 0.4 mg/mL proteinase (Sigma Type XXIV, 7–14 U/mg). After 45 minutes, the digestion medium was replaced with fresh Ca²⁺-free buffer containing collagenase only, and the suspension was gently triturated with a Pasteur pipette for 5–20 minutes. Enzymatic activity was stopped by adding 2% bovine calf serum. The cell-containing supernatant was collected and centrifuged (58 g, 5 min).

For further processing, the residual tissue was resuspended in storage solution containing (mM): taurine 20, glutamic acid 70, KCl 30, KH₂PO₄ 10, MgCl₂ 1, HEPES 10, BDM 10, glucose 11, and 2% bovine calf serum (pH 7.4, adjusted with KOH, 37 °C). Additional cardiomyocytes were released by gentle pipetting with a serological pipette. The suspension was centrifuged again, and cell pellets were resuspended in storage solution. Only preparations enriched in elongated cardiomyocytes with visible cross-striations were used. These cells were plated onto laminin-coated recording chambers and allowed to settle for 30 minutes before experiments, which were performed at room temperature.

### Action potential recordings

Action potentials were measured using the whole-cell patch-clamp method in a current-clamp mode. Recording pipettes (3–5 MΩ) were filled with an internal solution containing (mM): 92 K-aspartate, 48 KCl, 1 Mg-ATP, 10 HEPES, 0.02 EGTA, 0.1 GTP-Tris, and 4 Na₂-ATP (pH adjusted to 7.2 with KOH). The external bath solution consisted of (mM): 140 NaCl, 4 KCl, 1 MgCl₂, 2 CaCl₂, 10 glucose, and 10 HEPES (pH adjusted to 7.4 with NaOH). To trigger action potentials, square current pulses (1–2 nA, 1–5 ms duration) were applied at a frequency of 1 Hz. After achieving whole-cell access, series resistance was typically 5–15 MΩ. Capacitance transients were compensated in the cell-attached state, and both membrane capacitance and series resistance were corrected following rupture. Signals were acquired using an EPC10 amplifier (HEKA Elektronik) and filtered with Bessel filters at cutoff frequencies of 2.9 and 10 kHz.

### Differentiation and maintenance of iPSC-CMs and iPSC-CFs

Human iPSC-aCMs were purchased from Axol Bioscience (cat. no. ax2508). Cells were thawed and seeded on fibronectin-coated culture vessels and were maintained in cardiomyocyte maintenance basal medium (Axol Bioscience, cat. no. ax2530) as per the manufacturer’s instructions.

Human iPSC-vCMs were differentiated from a human iPSC line BJ RiPS-D, which was derived from fibroblasts of a healthy Caucasian male donor^56^, using a modified protocol from Limpitikul et al^57^. Briefly, iPSCs were cultured in Geltrex matrix (Thermo Scientific, cat. no. A1413302)-coated tissue culture plates and fed daily with Essential 8 medium (Thermo Scientific, cat. no. A1517001). When ∼ 95% confluent (day 0), medium was changed to RPMI-1640 GlutaMAX medium (Thermo Scientific, cat. no. 61870127) supplemented with B-27 (minus insulin) (Thermo Scientific, cat. no. A1895601) and 6 µM CHIR99021 (Stemgent, cat. no. 04-0004). Cells were maintained in RPMI-1640 GlutaMAX medium supplemented with B-27 (minus insulin) for the first 7 days, with medium exchange every 2 days. On day 3, 5 µM IWP-4 (Stemgent, cat. no. 04-0036) was added. On day 8, medium was changed to RPMI-1640 GlutaMAX with B-27 supplement (Thermo Scientific, cat. no. 17504044), with medium exchange every 2 days. Spontaneous contraction of iPSC-CMs was observed by days 8-9. IPSC-vCMs were then replated at half density on day 11 for metabolic selection^58^ on days 15-20. On day 25, cells were replated again on Geltrex matrix–coated optically clear imaging 96-well plates at 5×10^4^ cells/well. On optically clear imaging plates to create monolayers for imaging. IPSC-vCMs were used for functional studies on days 32-35. Detailed characterization of iPSC-aCMs and iPSC-vCMs is shown in **Supplementary Fig. 9**.

Human iPSC-CFs were differentiated from the BJ RiPS-D cell line using a previously published protocol^59^. Briefly, when iPSCs reach ∼ 95% confluence (day 0), the medium is exchanged with RPMI-1640 GlutaMAX supplemented with B-27 (minus insulin) and 6 µM CHIR99021. Cells were maintained in RPMI-1640 GlutaMAX medium supplemented with B-27 (minus insulin) for the first 7 days, with medium exchange every 2 days. On day 3, 5 µM IWR-1 (Selleckchem, cat. no. S7086) was added. On day 6, cells were replated at 2×10^5^ cells/cm^2^ density in Advanced DMEM/GlutaMAX medium (Life Technologies, cat. no. 12634-028) containing 5 μM of CHIR99021, 2 μM of retinoic acid (Sigma, cat. no. R2625), 5 μM of Y27632, and 1% fetal bovine serum. Cells were maintained in Advanced DMEM/GlutaMAX from days 6-13 with medium change every 2 days. On days 7-8, 5 μM of CHIR99021 and 2 μM of retinoic acid were added. On day 11, cells are replated at the 2×10^5^ cells/cm^2^ density in Advanced DMEM/GlutaMAX supplemented with 2 μM SB431542 (Tocris Bioscience, cat. no. 1614). At this step, a confluent sheet of proepicardial cells was achieved. On days 14-19, proepicardial cells were cultured in fibroblast growth medium 3 (PromoCell, cat. no. C23025) containing 20 ng/ml of FGF2 (R&D Systems, cat. no. 233-FB) and 10 μM of SB431542. On day 20 and every other day thereafter, cardiac fibroblasts were fed with fibroblast growth medium 3, supplemented with 10 μM of SB431542. IPSC-CFs were seeded at 1×10^5^ cells/cm^2^ for usage. Detailed characterization of iPSC-CFs and their epicardial precursors is shown in **Supplementary Fig. 10**.

### Differentiation of human macrophages

Human monocyte cell line, THP-1, was differentiated into macrophages based on a modified protocol from Phu et al^60^. Briefly, THP-1 cells were cultured in suspension in RPMI-1640 medium (Thermo Scientific, cat. no. 11875093), supplemented with 10% fetal bovine serum, 1% GlutaMAX (Thermo Scientific, cat. no. 35050061), and 0.05 mM β-mercaptoethanol. When confluent, cells are then seeded on tissue culture-treated plates at a density of 2.8×10^5^ cells/cm^2^ and differentiated into macrophages by culturing in 25 ng/mL phorbol 12-myristate 13-acetate (PMA, Sigma-Aldrich, cat.no. P1585) for 48 hours. Differentiated macrophages are then switched to AIM V serum-free medium (Thermo Scientific, cat. no. 12055083) for at least 48 hours before usage.

### Isolation of and treatment with VAT EVs

VAT biopsies were obtained from donors as part of the EVOC study (EVs in Obesity and Cardiometabolic disease, NCT06408961). Routinely discarded visceral fat was collected during the sleeve gastrectomy surgery, and in gastric bypass, where no visceral fat is routinely removed, a small segment of visceral fat was removed from the greater omentum. VAT was minced and cultured in RPMI 1640 media (Thermo Scientific, cat. no. 11875093) with B27 supplement (Thermo Scientific, cat. no. 17504044). Conditioned media, containing EVs released from VAT, were collected after 48 hours and 96 hours, centrifuged and filtered (0.8 um) to remove tissue debris, and stored at -80 °C until usage.

Small EVs were isolated from VAT conditioned media by size exclusion chromatography (SEC)-based method using qEVoriginal 35 nm Gen 2 columns (Izon Science, cat. no. ICO-35). The void volume was set at 2.5 mL, and 3 mL of EV-enriched fractions in phosphate buffered saline (PBS; Thermo Scientific, cat. no. 10010049) were collected. Concentration and size distribution of EVs were measured by microfluidic resistive pulse sensing (Spectradyne’s nCS1, Signal Hill). Cells were treated with 7.5 x 10^10^ EVs/mL for 24 hours for iPSC-CMs, 96 hours for iPSC-CFs (re-dosed at the same concentration after 48 hours), and 48 hours for macrophages. As controls, iPSC-CMs, iPSC-CFs, and macrophages were treated with an equal number of EVs extracted from conditioned medium from respective cell types for the same duration.

### Measurement of electrical properties and calcium transients of iPSC-CMs

For action potential measurement, iPSC-CMs were stained with voltage-sensitive fluorescence dye, FluoVolt (Thermo Scientific, cat. no. F10488) for 20 minutes at 37 °C, following the manufacturer’s instructions, washed with Tyrode’s solution three times, and imaged in Tyrode’s solution. For calcium transient measurement, iPSC-CMs were stained with calcium-sensitive fluorescence dye, Fluo-4 NW (Thermo Scientific, cat. no. F36206) for 30 minutes at 37 °C, following the manufacturer’s instructions, washed three times, and imaged in Tyrode’s solution. Both action potential and calcium transient measurements were performed using an automated video imaging system, Kinetic Imaging Cytometry (Vala Sciences), at 95 frames per second. Image intensity was measured using ImageJ^61^. Custom software written in MATLAB (Mathworks) was used to calculate action potential duration, calcium transient magnitude, time to peak, and decay constant.

### Assessment of gene expression with quantitative real-time PCR

Total RNA was isolated using miRNeasy Micro Kit (Qiagen, cat. no. 217084). RNA concentration was measured using a NanoDrop (Thermo Scientific). Equal amount of total RNA (250 – 1000 ng) was used to create cDNA libraries using a high-capacity cDNA reverse transcription kit (Thermo Scientific, cat. no. 4368813). Quantitative real-time PCR was performed using Ssoadvanced Universal SYBR (Bio-Rad, cat. no. 1725274) in the QuantStudio 6 Flex real-time PCR system (Thermo Scientific). Primers used are listed in **Supplementary Table 11**. For multiple hypothesis testing for genes of interest, the Bonferroni correction was applied. Statistical significance was set at p < 0.05.

### Quantification of collagen secretion by iPSC-derived cardiac fibroblasts

Medium from EV-treated iPSC-CFs was collected at 48 and 96 hours and combined. Secreted collagen in medium was then measured using Sirius Red Collagen Detection Kit (Amsbio, cat. no. 9062), following the manufacturer’s instructions.

### Animal studies

Exomap mice were a kind gift from Dr. Stephen Gould at Johns Hopkins University^26^. Adipoq-Cre BAC transgenic mice, where Cre recombinase is expressed under the control of adiponectin promoter/enhancer region, were obtained from The Jackson Laboratory (strain no. 028020). Transgenic mice were crossed with wild-type (C57BL/6J, The Jackson Laboratory) to produce litters containing both transgenic and wild-type offspring. Genotyping was performed by PCR on DNA extracted from ear tissue, using the following primer set: forward 5’- GGTGATAGGTGGCAAGTGGTATTCCGTAAG -3’, reverse 5’- CATA-TATGGGCTATGAACTAATGACCCCGT -3’ for Exomap and forward 5’- GGATGTGCCATGTGAGTCTG -3’, reverse 5’- ACGGACAGAAGCATTTTCCA -3’ for Adipoq-Cre. Starting at 8 weeks of age, the double positive Adipoq-Cre^+^/Exomap^+^ male mice were fed with either high-fat diet (Research Diets, cat. no. D12492, 60% fat by calories) for obese mice or regular chow for lean mice. After 16 weeks of diet, the mice were sacrificed, and their hearts were collected.

### Flow cytometry of cardiac macrophages

Macrophages were isolated from left ventricular tissue using a protocol described in Hulsmans et al^62^. Briefly, mice hearts were perfused with cold PBS. Left ventricles were cut out, minced, and incubated in enzyme cocktails at 37 °C for 40 minutes while on a shaker at 200 rpm. Enzyme cocktails include 450 U/ml of collagenase I (Sigma-Aldrich, cat. no. C0130), 125 U/ml of collagenase XI (Sigma-Aldrich, cat. no. C7657), 60 U/ml of DNase I (Sigma-Aldrich, cat. no. D5319), and 60 U/ml of hyaluronidase (Sigma-Aldrich, cat. no. H3506) and 20 mM HEPES. Cells were strained through a 40-μm cell strainer, centrifuged at 340 xg for 5 minutes, and resuspended in PBS.

Cells were stained with the following antibodies at 4 °C for 30 minutes (company, catalog no, dilution): APC-Cy7-conjugated anti mouse CD45 (Biolegend, 103115, 1:50), BV650-conjugated anti mouse/human CD11b (Biolegend, 101239, 1:25), and BV785-conjugated anti mouse CD206 (Biolegend, 141729, 1:50). 7-AAD viability staining solution (Biolegend, 420403, 1:16) was used as viability marker. Then stained cells were washed with MACS buffer (0.5% bovine serum albumin and 2 mM EDTA in Dulbecco’s Phosphate Buffered Saline), centrifuged at 340 xg for 5 minutes, and resuspend in MACS buffer for flow cytometry. Data were acquired on an FACSAriaIIu (BD Biosciences) and analyzed with FlowJo software (FlowJo, LLC). Macrophages were identified as CD45^+^/CD11b^+^. M1 macrophages were CD206^-^ and M2 macrophages were CD206^+^.

### Imaging of adipose tissue

After the hearts were harvested, inguinal white adipose tissue (iWAT) was dissected from the mice. A small 3 x 3 x 3 mm piece of iWAT was placed on microscope slide and a coverslip was pressed on top of the iWAT to flatten the tissue for imaging. Prepared slides of flattened iWAT were imaged under fluorescence microscope using standard green and red fluorescence cubes.

### Immunoblotting

Western blot analysis was performed on snap-frozen left atrial and left ventricular tissue. Tissue samples were lysed in Pierce^TM^ RIPA lysis and extraction buffer (Thermo Scientific, cat. no. 89900) supplemented with Halt^TM^ protease and phosphatase inhibitor cocktail (Thermo Fischer Scientific, cat. no. 78441). Protein concentration was measured using Pierce^TM^ BCA protein assay kits (Thermo Scientific, cat. no. 23227). Equal amounts of protein lysate (23 μg) were treated with Laemmli buffer with β-mercaptoethanol and incubated at 95 °C for 7 minutes. The lysate was then electrophoresed on a 4-20% SHS-polyacrylamide resolving gel and transferred to a PVDF membrane. Membranes were incubated overnight at 4 °C with primary antibodies, washed, and incubated for 1 hour at room temperature with secondary antibodies. Primary antibodies used include (company, catalog no, dilution): mNeonGreen (Chromotek, 32f6-100, 1:1000), vinculin (Sigma-Aldrich, V9264, 1:10,000). Secondary antibodies include (company, catalog no, dilution): HRP-conjugated goat anti-rabbit immunoglobulins (Agilent Technologies, P044801-2, 1:2500) and horseradish peroxidase (HRP)-conjugated goat anti-mouse immunoglobulins (Agilent Technologies, P044701-2, 1:5000). Lastly, antibody signal was detected using Supersignal^®^ West Femto Maximum Sensitivity (Thermo Scientific, cat. no. 34095). The intensity of proteins of interest was quantified using iBright Analysis Software (Thermo Fisher Scientific).

### Nano-flow cytometry analysis of circulating EVs

Mouse whole blood was collected by buccal puncture into EDTA-containing tubes and centrifuged at 2000 xg for 10 minutes at 4 °C. Supernatant (plasma) was then collected. Small EVs were isolated from plasma by size exclusion chromatography (SEC)-based method using qEVoriginal 35 nm Gen 2 columns (Izon Science, cat. no. ICO-35). The void volume was set at 2.5 mL, and 3 mL of EV-enriched in PBS were collected. Isolated EVs were concentrated back down to 0.5 mL using Amicon® Ultra Centrifugal Filter, 10 kDa MWCO (Millipore, cat. no. UCF8010).

Concentration of isolated plasma EVs was measured using nano flow cytometry was performed using a flow nanoanalyzer (NanoFCM). An aliquot of 1-2×1011 EVs/mL was diluted 10-fold with PBS and fixed with permeabilization buffer (NanoFcm, cat. no. P2320) at 1:5 ratio for 30 minutes at room temperature. Next, fixed sample was diluted 1:10 fold with PBS and incubated at room temperature with mNeonGreen primary antibody (Chromotek, 32f6-100) at 1:20 concentration for 30 minutes. Next, secondary antibody, Goat anti-Mouse IgG (H+L) Cross-Adsorbed, Alexa Fluor™ 488 (Invitrogen, cat. no. A-11001) was added to the mixture at 1:1000 concentration and the mixture was incubated at room temperature for 30 minutes. Samples were diluted at 1:100 – 1:200 with PBS-H (10mM HEPES in PBS, pH 7.4) and analyzed with a flow nanoanalyzer using a standard green fluorescence filter.

### Immunohistochemistry

Mice hearts were perfused with cold PBS, dissected, and fixed overnight at 4 °C with 4% paraformaldehyde. Fixed hearts were transferred to 70% ethyl alcohol and were paraffin-embedded. Thin sections of tissue were cut, underwent deparaffinization, antigen retrieval, permeabilized with 0.5% Triton® X-10 (Thermo Scientific, cat. no. PI85111) in PBS, and blocked with 3% bovine serum albumin in PBS prior to incubation with primary antibodies overnight at 4 °C, and later incubation with secondary antibodies for 1 hour at room temperature. Tissue sections were finally counterstained with DAPI. Primary antibodies used include (company, catalog no, dilution): mNeoGreen (Chromotek, 32F6, 1:50), cardiac troponin I (Abcam, ab47003, 1:100), vimentin (Abcam, ab92547, 1:100), and CD68 (Cell Signaling, 97778T, 1:200). Secondary antibodies include (company, catalog no, dilution): Goat anti-Mouse IgG (H+L) Cross-Adsorbed Secondary Antibody, Cyanine3 (Invitrogen, A10521, 1:500) and Goat anti-Rabbit IgG (H+L) Cross-Adsorbed Secondary Antibody, Cyanine5 (Invitrogen, A10523, 1:500).

### Bulk RNA sequencing and analysis

Total RNA was isolated using miRNeasy Micro Kit (Qiagen, cat. no. 217084). Samples were treated with DNase I and cleaned up using RNA Clean & Concentrator-5 (Zymo Research, cat. no. R1014). Stranded cDNA libraries were created using SMARTer Stranded Total RNA-Seq Kit v2 - Pico Input Mammalian (Takara Bio USA, Inc., cat. no. 634413) and sequenced at paired-end 150 bp on the Illumina NovaSeq Xp platform (Illumina).

Adapter sequences were trimmed from sequencing reads using Cutadapt (v4.8)^63^. Next, Quality assessment of both raw and adaptor-trimmed reads was carried out with FastQC (v0.12.1) (www.bioinformatics.babraham.ac.uk/projects/fastqc). Subsequently, the trimmed reads were aligned to the Gencode GRCh38.p13 reference genome using STAR (v2.7.11a)^64^. STAR was supplied with Gencode v38 gene annotations to enhance mapping precision, and gene-level read counts were obtained using featureCounts (v2.0.6)^65^. ComplexHeatmap^66^ was used for cluster analysis and visualization. Significantly differentially expressed genes with absolute fold change ≥ 1.2 and FDR p-value ≤ 0.1 were detected by DESeq2 (v1.42.1)^67^. For differential expression analyses, genes with a median read count below 5 in both conditions were excluded. To account for the batch effects arising from the sequencing of active fibroblast and macrophage samples across two distinct batches, two batch correction methods were employed for differential expression analysis. First, DESeq2 was utilized with batch included as a covariate in the model. Second, DESeq2 was applied to data pre-processed using ComBatSeq^68^ to remove batch effects. Only genes identified as differentially expressed by both approaches were reported to ensure robust and reliable results. Genome Ontology and KEGG pathway over-representation analysis (ORA) was performed on differentially expressed genes using the WebGestaltR package (v1.0.0)^69^. Gene set enrichment analysis (GSEA) was performed using the GSEA package (v4.3.2)^70^ on the database (v2022.1.Hs). For both ORA and GSEA, differential expression results derived from ComBat-Seq pre-processed data were utilized for the active fibroblast and macrophage datasets.

### Transcriptome-wide genetic association studies (TWAS) and localization of cell types of interest via epigenetics and chromatin-based studies

Using summary statistics from genome-wide association studies (GWAS) of QT interval (n=117,532)^71^ and atrial fibrillation (n=1,650,345 with 181,446 cases)^72^, we performed transcriptome-wide association studies (TWAS)^73^ of the corresponding phenotypes. We used MR-JTI^74^ TWAS models trained on two heart tissues (left ventricle and atrial appendage), and subset the associations to retain only genes that were differentially expressed (FDR-adjusted p-value < 0.05) upon EV treatment in atrium, ventricle, fibroblast, or monocyte. Since each subset of TWAS genes consisted of ≈900 genes, TWAS associations with *p* < 5.56 x 10^-5^ (Bonferroni adjusted for 900 comparisons) were considered significant.

Since TWAS was performed in atrial and ventricular tissue, inference on whether specific genes in a specific cell type (e.g., atrial and ventricular cardiomyocyte, fibroblasts, macrophages) are related to our phenotypes is inaccessible to standard TWAS methodology. Therefore, we utilized cell-type-specific epigenomic, chromatin accessibility, and chromatin conformation assays to provide <u>cell-type-specific</u> genetic evidence linking QT interval and AF with the VAT-EV induced gene expression changes in the four cell types tested. Our methodology builds on previous work^21^, with attribution provided by this statement. We utilized a regulatory annotation framework that integrates 18 epigenomic features, including proximity to a transcription start site, assays of chromatin accessibility (ATAC-seq and DNase-seq), histone ChIP-seq (for chromatin marks H3K27ac, H3K27me3, H3K4me1, H3K4me2, H3K4me3, H3K36me3, H3K79me2, H3K9ac, H3K9me3, and H4K20me1), and transcription factor ChIP-seq (for RNA polymerase II subunit POL2RA and cohesion complex subunits RAD21 and SMC3)^21,75^. This framework was used to identify cell type-specific regulatory elements in four atrium biosamples, eight ventricle biosamples, 41 fibroblast biosamples, and four macrophage-analog biosamples^76^. We utilized Hi-C data from the 4D Nucleome Project^77^ to identify chromatin contacts between regulatory elements and differentially expressed genes. Hi-C in primary cardiomyocytes from the left atrium (4DNFIBNXTSV4) was used to represent chromatin contacts in atrium, Hi-C in primary cardiomyocytes of the left ventricle (4DNFISWHXA16) was used to represent chromatin contacts in ventricle, Hi-C in the cell line IMR-90 (4DNFIJTOIGOI) was used to identify chromatin contacts in fibroblasts, and Hi-C in monocytes (4DNFI4HZHGLC) was used for chromatin contact analysis in macrophages. For each Hi-C dataset, the contact matrix had been processed using the gold standard 4D Nucleome pipeline (https://data.4dnucleome.org/resources/data-analysis/hi_c-processing-pipeline) and normalization had been performed using the iterative correction and eigenvalue decomposition (ICE) algorithm^78^. Cooler (v0.8.2)^79^, was downloaded in mcool format. Contact pairs between a regulatory element and one of the differentially expressed genes were then exported in bedpe format. Using BEDTools (v2.30.0)^80^, we identified genome-wide significant (*p* < 5 x 10^-8^) GWAS SNPs associated with QT interval^71^ and AF^72^ that overlapped one of the cell-type-specific regulatory elements in contact (via Hi-C) with a differentially expressed gene.

### TRPC3 suppression

IPSC-aCMs and iPSC-vCMs were treated with Pyr3 (Tocris, cat. no. 3751), a small molecule TRPC3 inhibitor, at 7.5 μM for 24 hours at the same time as VAT EV treatment. Myocytes were also treated with the same concentration of Pyr3 during action potential recording.

## Acknowledgements

We thank Dr. Kyle Metkowski for his advice on flow cytometry and Dr. Guoping Li on differentiation of iPSC-vCMs. We acknowledge all the participants in the human atrial tissue study from the University of Regensburg and the University of Göttingen, participants in the EVOC study, and the participants in the Mass General Brigham Biobank.

## Funding

This work was supported by a grant from AHA (20SFRN35120267) and NIH (NHLBI 1R35HL150807) to Dr. Das and AHA (20SFRN35120267) to Dr. Shah. Dr. Gamazon was supported by NIDDK U01DK140952 and a grant from the Vanderbilt Diabetes Center (Functional Multi-Omics of Diabetes and Metabolism Core). Dr. Limpitikul received support from John S. LaDue Memorial Fellowship in Cardiovascular Medicine from Harvard medical school.

## Declaration of interests

Dr. Das is a founding member, owns equity and consults for Thryv Therapeutics and Switch Therapeutics that did not play any role in this study and is a consultant for Renovacor. Dr. Shah is supported by grants from the National Institutes of Health. Dr. Shah has equity ownership in and is a consultant for Thryv Therapeutics. Dr. Shah is a co-inventor on pending patents or disclosures on molecular biomarkers of fitness, lung disease, cardiovascular diseases and phenotypes, and metabolic health, use of RNAs (including spatial) as therapeutics and diagnostic biomarkers in disease, and methods in metabolomics. Dr. Gamazon has performed consulting for Thryv Therapeutics and has pending patents or disclosures on molecular biomarkers of fitness, cardiovascular diseases and phenotypes, and metabolic health, use of RNAs as therapeutics and diagnostic biomarkers in disease, and methods in metabolomics. Dr. Pabel is employed by the Novartis Institute for Biomedical Research.

## Data availability

Data has been deposited in the NIH GEO repository (pending accession number).

## Notes

### Competing Interest Statement

The authors have declared no competing interest.

