## Supplementary Information for "A role of extracellular vesicle-mediated inter-organ communication in obesity-related arrhythmia"

### SUPPLEMENTARY METHODS

#### Immunoblotting of EVs

Western blot analysis was performed on VAT EV and plasma-derived EV. EVs were lyzed with 10x cell lysis buffer (Cell Signal Technology, cat. no. 9803S) supplemented with Halt™ protease and phosphatase inhibitor cocktail (Thermo Fischer Scientific, cat. no. 78441). Protein concentration was measured using Pierce™ BCA protein assay kits (Thermo Scientific, cat. no. 23227). Equal amounts of protein lysate (23 μg) were treated with Laemmli buffer with and without β-mercaptoethanol. Protein electrophoresis, membrane blotting, and antibody incubation are described in the method section of the main text. Primary antibodies used include (company, catalog no, dilution): CD63 (Novus Biological, NBP2-42225, 1:1000), CD9 (System Biosciences, EXOAB-CD9A-1, 1:1000), alix (Abcam, ab88388, 1:2000), adiponectin (Cell Signaling, 2789, 1:1000). Samples probed with anti-CD63 and anti-CD9 antibodies were run in a non-reducing condition (without β-mercaptoethanol).

#### Lactate dehydrogenase cytotoxicity assay

Medium was collected after 24 hours of EV treatment, centrifuged at 500 xg for 5 min and then 2000 xg for 10 minutes to remove cell debris. LDH Cytotoxicity Assay Kit (Cayman Chemical Company, cat. no. 601170) was used to measure released lactate dehydrogenase (LDH) released from cells as per the manufacturer's instructions.

#### Isolation of and treatment with plasma EVs

Plasma samples were obtained from the Mass General Brigham Biobank (IRB # 2009P002312) and donors from the EVOC study (EVs in Obesity and Cardiometabolic disease,

NCT06408961). Whole blood was first centrifuged at 2000 xg for 10 minutes. The clear yellow supernatant was collected and re-centrifuged at 3000 xg for 10 minutes. The supernatant (plasma) was collected and stored at -80 °C until usage.

Small EVs were isolated from 0.5 mL of plasma by size exclusion chromatography (SEC)-based method using qEVoriginal 35 nm Gen 2 columns (Izon Science, cat. no. ICO-35). The void volume was set at 2.5 mL, and 3 mL of EV-enriched fractions in phosphate buffered saline (PBS; Thermo Scientific, cat. no. 10010049) were collected. Isolated EVs were concentrated back down to 0.5 mL using Amicon® Ultra Centrifugal Filter, 10 kDa MWCO (Millipore, cat. no. UCF8010). For each 96-well, 75 uL of concentrated EVs and 75 uL of culture medium were added and cells for incubated for 24 hours.

**Diastolic and sarcoplasmic reticulum calcium measurements**

Myocytes were stained for 8 minutes at room temperature with a mixture of 2 µM fura-2 acetoxymethyl ester (fura-2, AM; Thermo Fisher, cat. no. F1201) and 0.02% pluronic acid. Cells were washed twice with Tyrode solution and incubation in Tyrode for additional 40 minutes was performed to allow de-esterification of fura-2, AM dye. Cells were then moved to Tyrode solution containing 2 mM calcium and 20 mM 2,3-butanedione monoxime.

Prior to calcium measurements, iPSC-aCMs and iPSC-vCMs were paced for 5-10 minutes at 1 Hz. Then, cells were treated with caffeine 10 mM for 5 s to estimate sarcoplasmic reticulum (SR) calcium content. Fluorescence intensity was measured at an emission wavelength of 510 nm with an excitation wavelength of 340 nm and 380 nm. The ratio of these two numbers ($F_{ratio}$) is used as an estimate for calcium concentration[1]. $F_{ratio}$ measured prior to addition of caffeine was used as an approximation of diastolic calcium concentration while peak $F_{ratio}$ after application of caffeine was used to estimate SR calcium content.

**Masson's Trichrome staining and collagen quantification of cardiac tissue**

Sectioned paraffin-embedded left ventricles were de-paraffinized. Masson's Trichrome staining was performed using VitroView™ Masson's Trichrome Stain Kit

(VitroVivo Biotech, cat. no. VB-3016) following the manufacturer's instructions.

Images of stained left ventricles were analyzed using colour_deconvolution2 plugin (https://github.com/landinig/IJ-Colour_Deconvolution2/tree/main) for the ImageJ software (National Institute of Health).

**Immunohistochemistry**

IPSC-aCMs, iPSC-vCMs, and iPSC-CFs were seeded on Geltrex-coated glass coverslips. Cells were fixed with 4% paraformaldehyde for 20 minutes at room temperature and washed 3 times with PBS. Cells were permeabilized with 0.2% Triton® X-100 (Thermo Scientific, cat. no. PI85111) in PBS for 10 minutes at room temperature and then blocked with 4% normal goat serum (Abcam, cat. no. ab7481) for 1 hour at room temperature. Cells were incubated with primary antibodies for 1 hour then washed 3 times with PBS. Next, cells were incubated with secondary antibodies for 1 hour at room temperature and washed 3 times with PBS. Coverslips were then mounted using ProLong™ Glass Antifade Mountant with NucBlue™ Stain with DAPI (Invitrogen, cat. no. P36981). Primary antibodies used include (company, catalog no, dilution): cardiac troponin I (Abcam, ab47003, 1:100), α-sarcomeric actinin (Abcam, ab9465, 1:200), myosin regulatory light chain 2, atrial isoform (MLC2a; Abcam, ab68086, 1:400), myosin light chain 2, ventricular isoform (MLC2v; Proteintech, 10906-1-AP, 1:200), Wilms tumor 1 (WT1; Abcam, ab89901, 1:300), α-smooth muscle actin (α-SMA; Abcam, ab7817, 1:200), collagen type 1 (COL1; Abcam, ab260043, 1:300). Secondary antibodies include (company, catalog no, dilution): Goat anti-Mouse IgG (H+L) Cross-Adsorbed Secondary Antibody, Alexa Fluor™ 635 (Invitrogen, cat. no. A-31574), Goat anti-Rabbit IgG (H+L) Cross-Adsorbed Secondary Antibody, Alexa Fluor™ 488 (Invitrogen, cat. no. A-11008), and Goat anti-Rabbit IgG (H+L) Highly Cross-Adsorbed Secondary Antibody, Alexa Fluor™ 635 (Invitrogen, cat. no. A-31577).

### SUPPLEMENTARY TABLES AND FIGURES

**Supplementary Table 1: Characteristics of right atrial tissue donor for action potential duration study**

| Characteristic | BMI > 25 (n=12 patients) | BMI < 25 (n=3 patients) | p-value |
|---|---|---|---|
| Body mass index (Mean±SD, $kg/m^2$) | 28.0±1.6 | 22.4±3.4 | 0.002 $_T$ |
| Male sex (%) | 83.3 | 66.7 | 0.52 $_F$ |
| Age (Mean±SD, y) | 66.8±8.8 | 66.0±7.8 | 0.90 $_T$ |
| Caucasian origin (%) | 100.0 | 100.0 | 0.99 $_F$ |
| Heart rate (Mean±SD, bpm) | 76.1±12.8 | 82.7±19.0 | 0.62 $_{MW}$ |
| Diabetes (%) | 25.0 | 33.3 | 0.99 $_F$ |
| Coronary artery disease (%) | 91.7 | 66.7 | 0.37 $_F$ |
| Ejection fraction (Mean±SD, %) | 51.5±11.9 | 56.0±9.0 | 0.54 $_{MW}$ |
| ACE inhibitor or ARB (%) | 83.3 | 50.0 | 0.40 $_F$ |
| B-Blocker (%) | 66.7 | 50.0 | 0.99 $_F$ |
| Aldosterone antagonist (%) | 33.3 | 0.0 | 0.99 $_F$ |
| Diuretic (%) | 50.0 | 100.0 | 0.47 $_F$ |
| Digitalis (%) | 8.3 | 0.0 | 0.99 $_F$ |
| Antiarrhythmic (%) | 0.0 | 0.0 | 0.99 $_F$ |
| Statins (%) | 83.3 | 50.0 | 0.40 $_F$ |

**Supplementary Table 2: Characteristics of visceral adipose tissue donors**

| Donor No. | age | sex | race | BMI | HTN | HLD | DM | Hgb A1C | NAFLD/NASH | AF | HFrEF |
|---|---|---|---|---|---|---|---|---|---|---|---|
| 1 | 39 | F | H | 42.0 | n | n | DM | 6 | n | n | n |
| 2 | 21 | F | B | 40.6 | n | n | pre-DM | 5.8 | n | n | n |
| 3 | 52 | F | W | 42.0 | y | n | DM | 8.7 | n | n | n |
| 4 | 59 | F | W | 42.1 | n | y | DM | 6.8 | n | n | n |
| 5 | 23 | M | W | 49.0 | n | n | n | 5.2 | n | n | n |
| 6 | 32 | F | W | 46.5 | n | n | pre-DM | 5.9 | n | n | n |
| 7 | 55 | F | B | 36.0 | y | y | DM | 8.2 | y | n | n |
| 8 | 54 | F | W | 36.6 | n | y | n | 5.4 | n | n | n |

y, yes; n, no; F, female, M, male; H, Hispanic; B, black/African American; W, white/Caucasian; BMI, body mass index ($kg/m^2$); HTN, hypertension; HLD, hyperlipidemia; DM, diabetes mellitus; pre-DM, pre-diabetes mellitus; Hgb A1C, hemoglobin A1C; NAFLD/NASH, non-alcoholic fatty liver disease/non-alcoholic steatohepatitis; AF, atrial fibrillation; HFrEF, heart failure with reduced ejection fraction.

**Supplementary Table 3: Characteristics of plasma donors**

**Lean plasma donors**

| Donor No. | age | sex | race | BMI | HTN | HLD | DM | Hgb A1C | NAFLD/NASH | AF | HFrEF |
|---|---|---|---|---|---|---|---|---|---|---|---|
| 1 | 70 | M | W | 24.4 | y | y | DM | 7.1 | n | n | n |
| 2 | 75 | F | W | 23.2 | y | y | n | 5.4 | n | n | n |
| 3 | 41 | M | W | 23.2 | n | y | n | 5.4 | n | n | n |
| 4 | 51 | F | W | 22.1 | n | n | n | 5.2 | n | n | n |
| 5 | 41 | F | W | 25.0 | n | n | n | na | n | n | n |
| 6 | 31 | F | W | 20.9 | n | n | n | 5.1 | n | n | n |

**Obese plasma donors**

| Donor No. | age | sex | race | BMI | HTN | HLD | DM | Hgb A1C | NAFLD/NASH | AF | HFrEF |
|---|---|---|---|---|---|---|---|---|---|---|---|
| 1 | 52 | M | W | 44.8 | y | y | n | 5.7 | n | n | n |
| 2 | 61 | F | W | 48.1 | y | y | n | 5.7 | n | n | n |
| 3 | 61 | M | W | 45.5 | y | n | n | 5.3 | n | n | n |
| 4 | 52 | F | W | 42.0 | y | n | DM | 8.7 | n | n | n |
| 5 | 32 | F | W | 46.5 | n | n | pre-DM | 5.9 | n | n | n |
| 6 | 55 | F | B | 36.0 | y | y | DM | 8.2 | y | n | n |

y, yes; n, no; F, female, M, male; H, Hispanic; B, black/African American; W, white/Caucasian; BMI, body mass index (kg/$m^2$); HTN, hypertension; HLD, hyperlipidemia; DM, diabetes mellitus; pre-DM, pre-diabetes mellitus; Hgb A1C, hemoglobin A1C; na, not available; NAFLD/NASH, non-alcoholic fatty liver disease/non-alcoholic steatohepatitis; AF, atrial fibrillation; HFrEF, heart failure with reduced ejection fraction.

**Supplementary Table 4: Differentially expressed genes in iPSC-aCMs upon VAT EV treatment**

**Supplementary Table 5: Differentially expressed genes in iPSC-vCMs upon VAT EV treatment**

**Supplementary Table 6: Overlapping differentially expressed genes between iPSC-aCMs and iPSC-vCMs upon VAT EV treatment**

**Supplementary Table 7: Differentially expressed genes in iPSC-CFs upon VAT EV treatment**

**Supplementary Table 8: Differentially expressed genes in macrophages upon VAT EV treatment**

**Supplementary Table 9: Significant TWAS associations for differentially expressed genes.**

**Supplementary Table 10: Significant TWAS associations for differentially expressed genes linked to a significant regulatory GWAS SNP.**

**Supplementary Table 11: Primers used in real-time PCR**

| Gene | Fwd Primers | Rev Primers |
|---|---|---|
| GAPDH | CAGCCTCAAGATCATCAGCAATG | CCATCCACAGTCTTCTGGGTG |
| 18s | GCAGAATCCACGCCAGTACAAG | GCTTGTTGTCCAGACCATTGGC |
| ACTB | CACCATTGGCAATGAGCGGTTC | AGGTCTTTGCGGATGTCCACGT |
| COL1A1 | GATTCCCTGGACCTAAAGGTGC | AGCCTCTCCATCTTTGCCAGCA |
| α-SMA | CTATGCCTCTGGACGCACAACT | CAGATCCAGACGCATGATGGCA |
| POSTN | CAGCAAACCACCTTCACGGATC | TTAAGGAGGCGCTGAACCATGC |
| CD11b | ACTTGCAGTGAGAACACGTATG | TCATCCGCCGAAAGTCATGTG |
| CD68 | GGAAATGCCACGGTTCATCCA | TGGGGTTCAGTACAGAGATGC |
| iNOS | TTCAGTATCACAACCTCAGCAAG | TGGACCTGCAAGTTAAAATCCC |
| TNFa | CCTCTCTCTAATCAGCCCTCTG | GAGGACCTGGGAGTAGATGAG |
| IL-1b | AGCTACGAATCTCCGACCAC | CGTTATCCCATGTGTCGAAGAA |
| IL6 | ACTCACCTCTTCAGAACGAATTG | CCATCTTTGGAAGGTTCAGGTTG |
| MCP1 | CAGCCAGATGCAATCAATGCC | TGGAATCCTGAACCCACTTCT |
| Arg1 | TGGACAGACTAGGAATTGGCA | CCAGTCCGTCAACATCAAAACT |
| TGFb1 | CTAATGGTGGAAACCCACAACG | TATCGCCAGGAATTGTTGCTG |
| IL10 | TCAAGGCGCATGTGAACTCC | GATGTCAAACTCACTCATGGCT |
| CD206 | TTGGCATTGCCTAGTAGCGTA | CTACAAGGGATCGGGTTTATGGA |

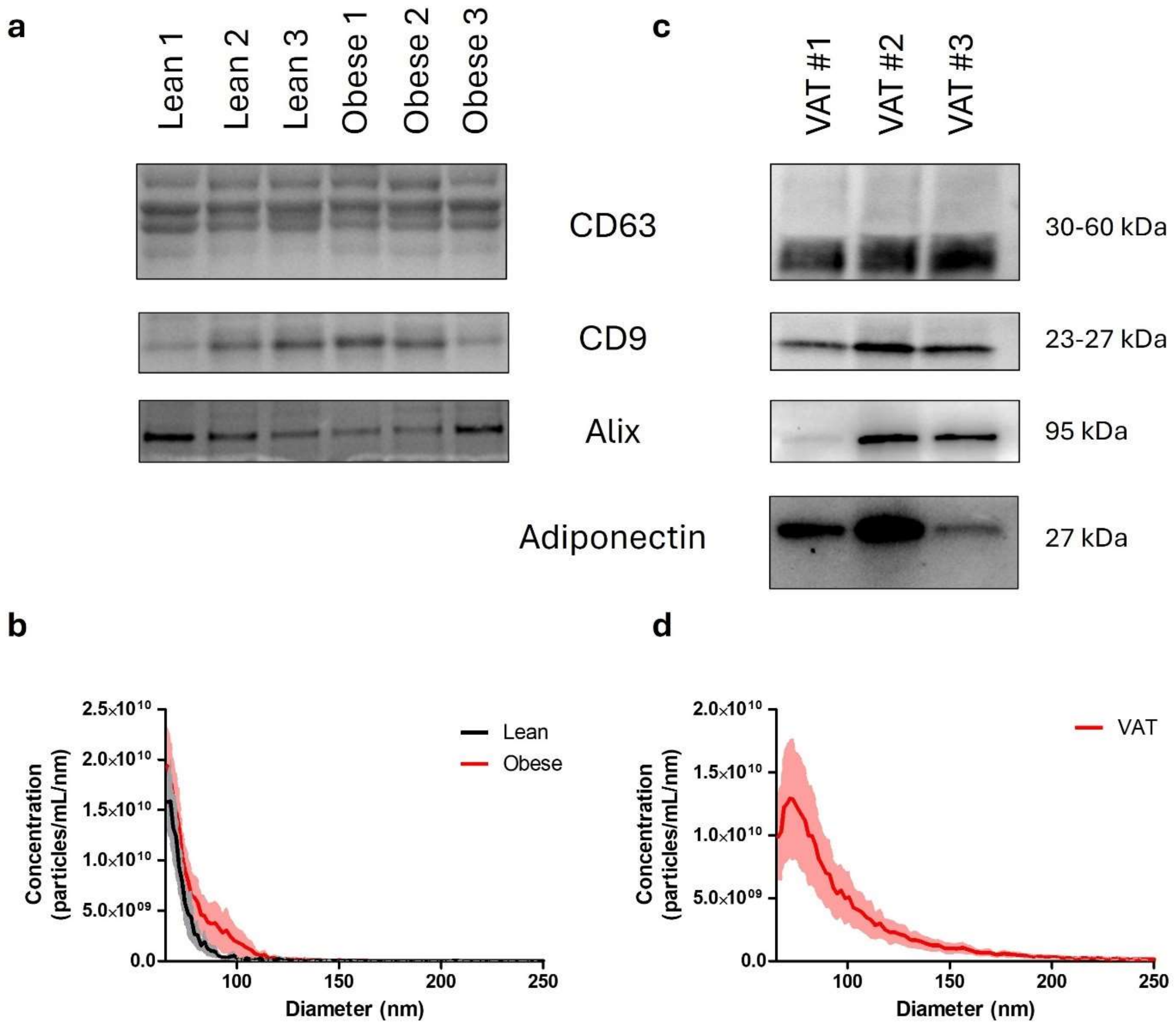


**Supplementary Figure 1: Characterization of plasma- and VAT-derived EVs. a,** Western blot of plasma-derived EVs from lean and obese donors shows the presence of canonical sEV markers, including tetraspanins (CD63, CD9) and cargo proteins (alix). **b,** Size distribution of plasma EVs by microfluidic resistive pulse sensing (MRPS). **c,** Western blot of VAT EVs demonstrates the presence of canonical sEV markers (CD63, CD9, alix) and adipose tissue marker (adiponectin). **d,** Size distribution of VAT EVs by microfluidic resistive pulse sensing (MRPS).

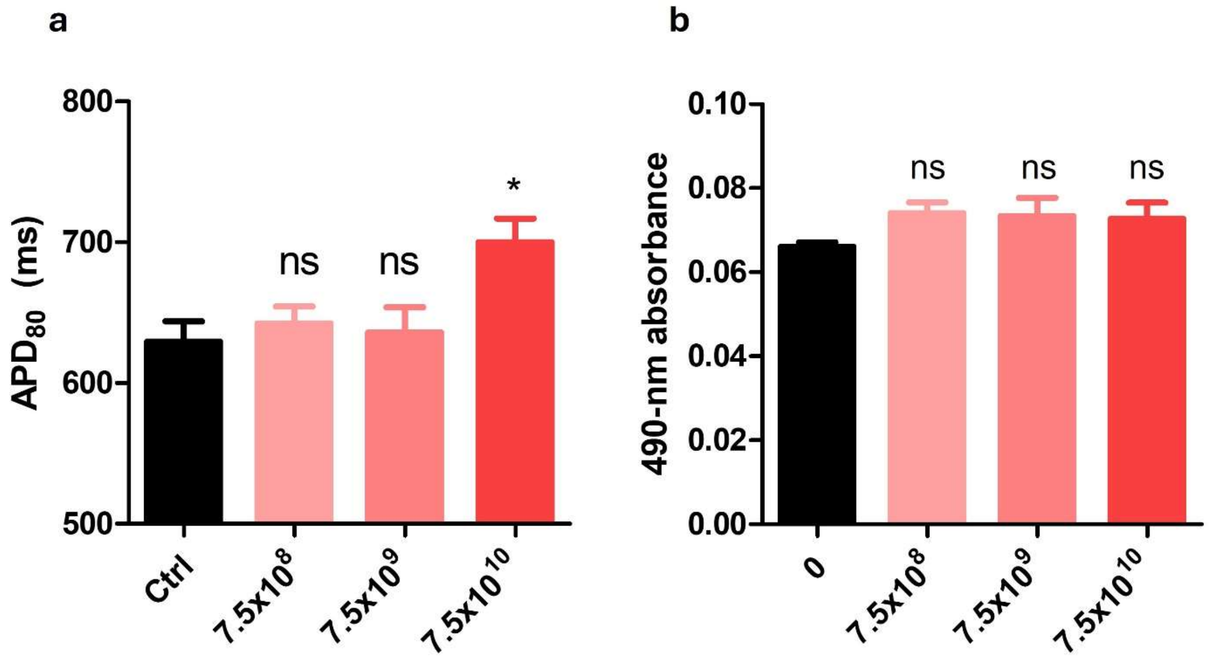


**Supplementary Figure 2: VAT EV dose response and cell toxicity. a,** IPSC-vCMs were treated for 24 hours with different concentration of EVs, including 7.5 x $10^8$ EVs/mL, 7.5 x $10^9$ EVs/mL, and 7.5 x $10^{10}$ EVs/mL prior to action potential measurement. Only iPSC-vCMs treated at 7.5 x $10^{10}$ EVs/mL show APD prolongation compared to control which were treated with iPSC-vCM-derived EVs at 7.5 x $10^{10}$ EVs/mL. **b,** Lactate dehydrogenase cytotoxicity assay shows no significant difference in level of LDH released by iPSC-vCMs treated with three different concentrations of VAT EVs versus untreated iPSC-vCMs.

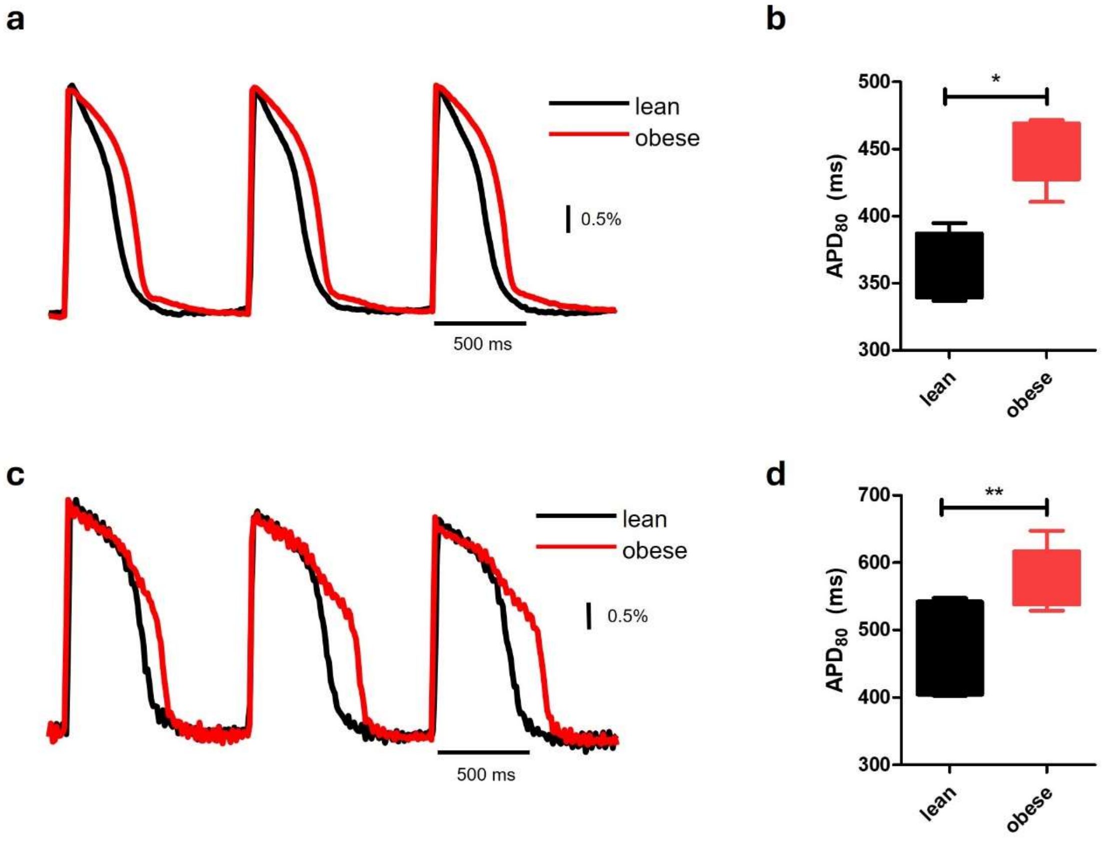


**Supplementary Figure 3: Treatment with obese plasma prolongs action potential duration. a,** Fluorescence signal representing change in membrane voltage of iPSC-aCMs during pacing at 1 Hz. Treatment of iPSC-aCMs with plasma-derived EVs from obese donors (red) prolongs APD compared to treatment with plasma-derived EVs from lean donors (black). **b,** IPSC-aCMs treated with plasma-derived EVs from obese donors have significantly prolonged APD compared to those treated with EVs from lean donors (n = 3 lean donors and 3 obese donors). APD was measured during pacing at 1 Hz. **c,** Representative membrane voltage recordings of iPSC-vCMs, paced at 1 Hz, showing APD prolongation upon treatment with plasma-derived EVs from obese donors (red) compared to lean donors (black). **d,** IPSC-vCMs treated with plasma-derived EVs from obese donors have significantly prolonged APD compared to those treated with EVs from lean donors (n = 3 lean donors and 3 obese donors). APD was measured during pacing at 1 Hz. *, $p < 0.05$, **, $p < 0.01$.

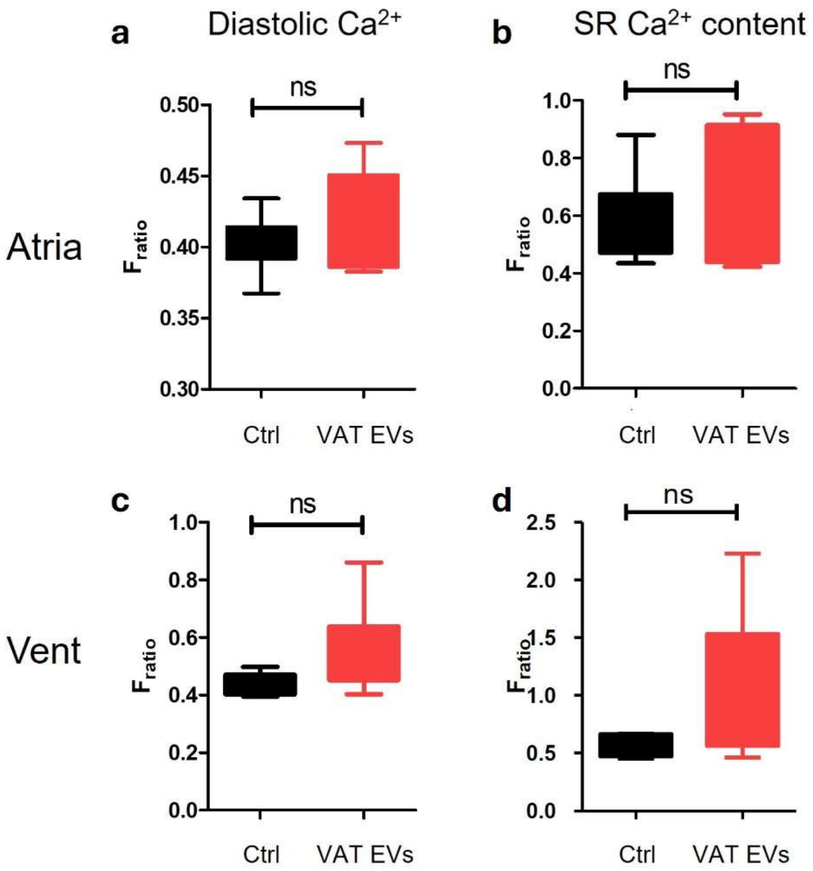


**Supplementary Figure 4: Diastolic $Ca^{2+}$ measurement and sarcoplasmic reticulum $Ca^{2+}$ load. a,** There is no significant difference in diastolic $Ca^{2+}$ level in iPSC-aCMs treated with VAT EVs (red, n = 6), compared to control (black, n = 7, treated with iPSC-aCM-derived EVs). **b,** There is no significant difference in sarcoplasmic reticulum (SR) $Ca^{2+}$ load between iPSC-aCMs treated with VAT EVs (red, n = 6) and control (black, n = 6, treated with iPSC-aCM-derived EVs). **c,** Similarly, there is no significant difference in diastolic $Ca^{2+}$ level between iPSC-vCMs treated with VAT EVs (red, n = 8) versus iPSC-vCM-derived EVs (black, n = 7). **d,** There is no significant difference between sarcoplasmic reticulum $Ca^{2+}$ content between iPSC-vCMs treated with VAT EVs (red, n = 5) versus iPSC-vCM-derived EVs (black, n = 6). $F_{ratio}$ is the ratio of fluorescence intensity at 340 nm to 380 nm emission.

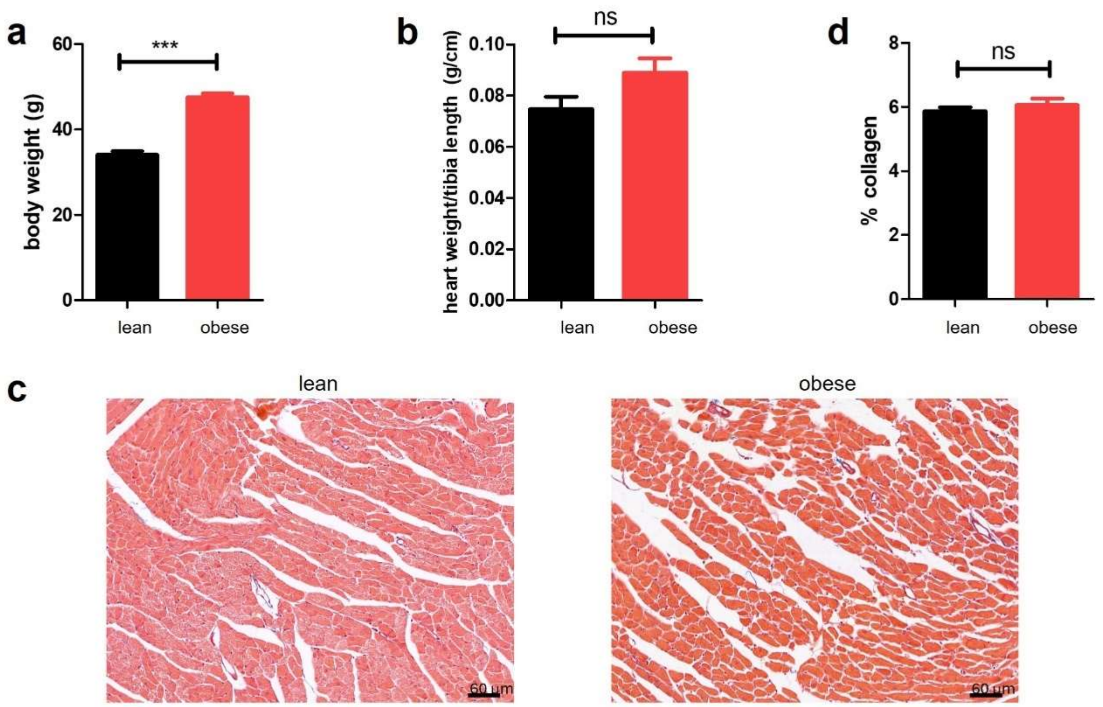


**Supplementary Figure 5: Characterization of the Adipoq-Cre$^+$/Exomap$^+$ model. a,** After 16 weeks of high fat diet, "obese" mice weigh significantly more than "lean" group which are on regular chow for 16 weeks (n = 5 lean, 5 obese, $p < 0.001$). **b,** There is no significant difference in heart weight normalized to tibial length in lean versus obese mice (n = 5 lean, 5 obese). **c,** Representative Trichrome staining of the left ventricles of lean and obese mice. **d,** Quantification of % collagen staining from Trichrome stain of the left ventricles of lean and obese mice shows no significant difference in the amount of fibrosis between the two groups (n = 13 randomly chosen images from 3 lean mice and 13 randomly chosen images from 3 obese mice).

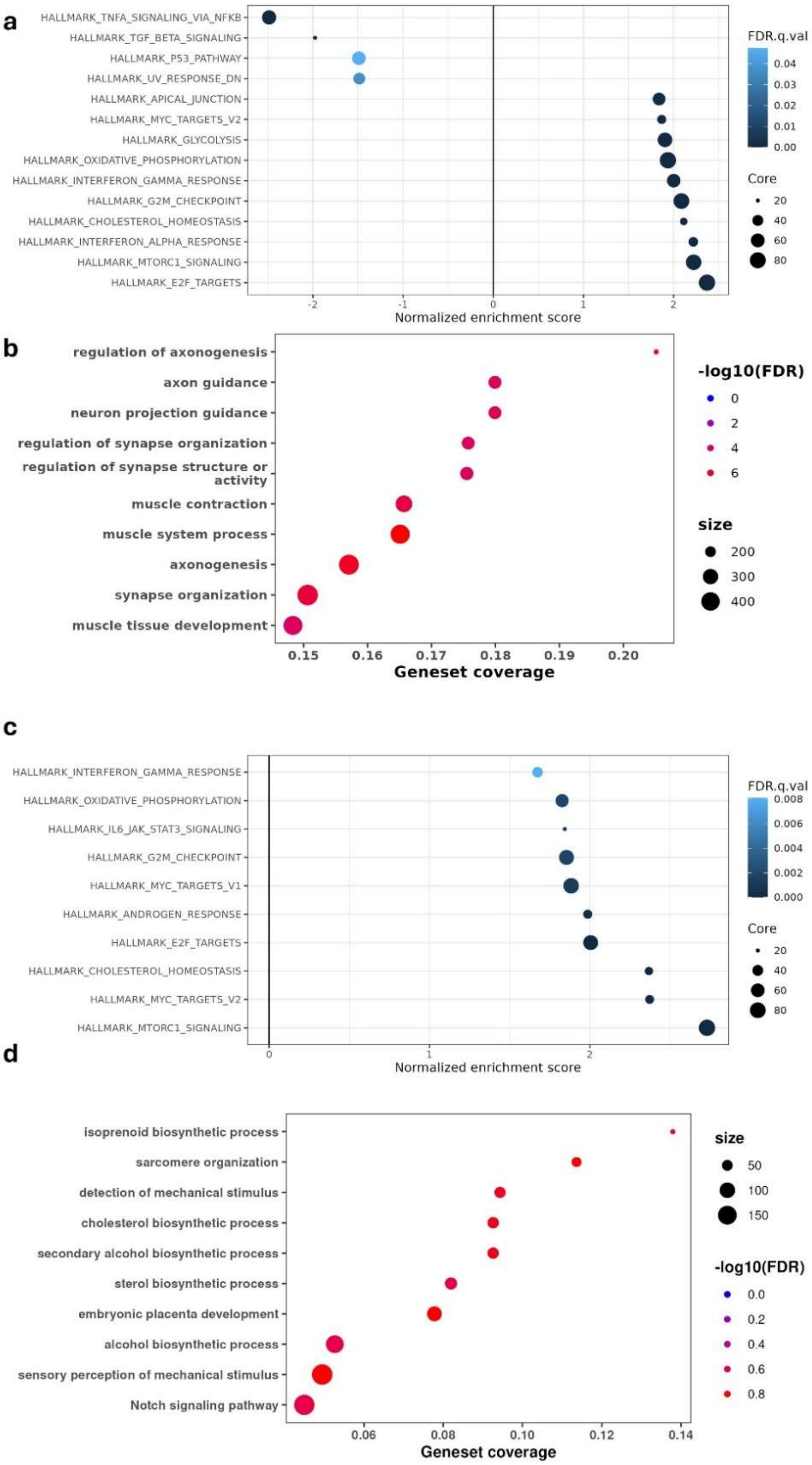

a
HALLMARK_TNFA_SIGNALING_VIA_NFKB
HALLMARK_TGF_BETA_SIGNALING
HALLMARK_P53_PATHWAY
HALLMARK_UV_RESPONSE_DN
HALLMARK_APICAL_JUNCTION
HALLMARK_MYC_TARGETS_V2
HALLMARK_GLYCOLYSIS
HALLMARK_OXIDATIVE_PHOSPHORYLATION
HALLMARK_INTERFERON_GAMMA_RESPONSE
HALLMARK_G2M_CHECKPOINT
HALLMARK_CHOLESTEROL_HOMEOSTASIS
HALLMARK_INTERFERON_ALPHA_RESPONSE
HALLMARK_MTORC1_SIGNALING
HALLMARK_E2F_TARGETS
-2
-1
0
1
2
Normalized enrichment score
FDR.q.val
0.04
0.03
0.02
0.01
0.00
Core
20
40
60
80
b
regulation of axonogenesis
axon guidance
neuron projection guidance
regulation of synapse organization
regulation of synapse structure or activity
muscle contraction
muscle system process
axonogenesis
synapse organization
muscle tissue development
0.15
0.16
0.17
0.18
0.19
0.20
Geneset coverage
-log10(FDR)
0
2
4
6
size
200
300
400
c
HALLMARK_INTERFERON_GAMMA_RESPONSE
HALLMARK_OXIDATIVE_PHOSPHORYLATION
HALLMARK_IL6_JAK_STAT3_SIGNALING
HALLMARK_G2M_CHECKPOINT
HALLMARK_MYC_TARGETS_V1
HALLMARK_ANDROGEN_RESPONSE
HALLMARK_E2F_TARGETS
HALLMARK_CHOLESTEROL_HOMEOSTASIS
HALLMARK_MYC_TARGETS_V2
HALLMARK_MTORC1_SIGNALING
0
1
2
Normalized enrichment score
FDR.q.val
0.008
0.006
0.004
0.002
0.000
Core
20
40
60
80
d
isoprenoid biosynthetic process
sarcomere organization
detection of mechanical stimulus
cholesterol biosynthetic process
secondary alcohol biosynthetic process
sterol biosynthetic process
embryonic placenta development
alcohol biosynthetic process
sensory perception of mechanical stimulus
Notch signaling pathway
0.06
0.08
0.10
0.12
0.14
Geneset coverage
size
50
100
150
-log10(FDR)
0.0
0.2
0.4
0.6
0.8

**Supplementary Figure 6: pathway enrichment analysis reveals distinct VAT EV-induced remodeling programs across cardiac cell types. a,** Hallmark pathway enrichment in iPSC-aCMs showing upregulation of proliferative pathways (MYC targets, E2F targets, G2M checkpoint), metabolic reprogramming (glycolysis, oxidative phosphorylation, cholesterol homeostasis), and stress responses (interferon signaling, mTORC1 signaling). Negative enrichment indicates downregulation of TGF-β signaling and DNA damage responses. Circle size represents core enrichment gene count, and color intensity indicates FDR-adjusted q-value significance. **b,** Gene Ontology biological process enrichment in iPSC-aCMs highlighting neuronal-like processes including axon guidance, synapse organization, and muscle contraction pathways, suggesting VAT EVs induce developmental reprogramming and altered excitation-contraction coupling. Circle size represents the number of genes in each pathway, color intensity indicates -log10(FDR), and geneset coverage shows the proportion of pathway genes that are differentially expressed. **c,** Hallmark pathway enrichment analysis showing normalized enrichment scores for significantly altered pathways in VAT EV-treated iPSC-vCMs compared to controls. VAT EVs upregulate metabolic and growth pathways including mTORC1 signaling, oxidative phosphorylation, MYC targets (V1 and V2), E2F targets, G2M checkpoint, and cholesterol homeostasis, indicating enhanced metabolic activity and cell cycle progression. Additional upregulated pathways include interferon gamma response, IL6-JAK-STAT3 signaling, suggesting inflammatory activation in ventricular cardiomyocytes. **d,** Gene Ontology biological process enrichment highlighting biosynthetic processes including isoprenoid, cholesterol, sterol, and alcohol biosynthesis pathways, sarcomere organization, detection of mechanical stimulus, embryonic placenta development, and Notch signaling pathway. The enrichment of biosynthetic pathways reflects enhanced metabolic activity, while sarcomere organization and mechanical stimulus detection suggest structural remodeling and altered mechanosensing in ventricular cardiomyocytes.

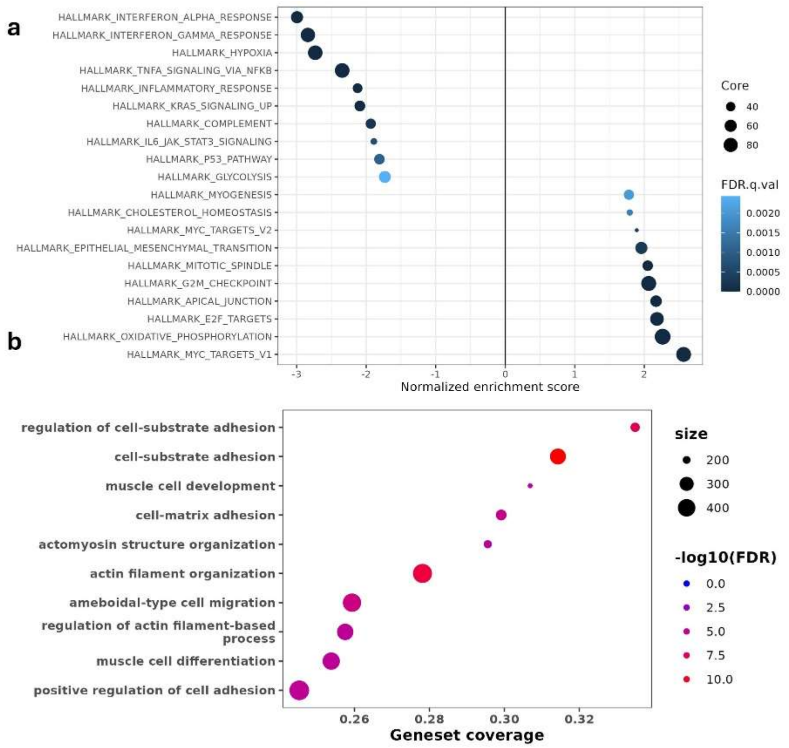


**Supplementary Figure 7: VAT EV treatment induces inflammatory activation and cytoskeletal remodeling in cardiac fibroblasts. a,**Hallmark pathway enrichment analysis showing normalized enrichment scores for significantly altered pathways in VAT EV-treated iPSC-CFs compared to controls. Proliferative and metabolic pathways (MYC targets, E2F targets, G2M checkpoint, oxidative phosphorylation, glycolysis) are upregulated. **b,** Gene Ontology biological process enrichment highlighting structural and adhesion-related processes including regulation of cell-substrate adhesion, cell-matrix adhesion, actomyosin structure organization, actin filament organization, and ameboidal-type cell migration. The enrichment of cytoskeletal organization pathways supports the acquisition of contractile properties characteristic of myofibroblast differentiation, while muscle cell development pathways reflect the transformation from quiescent fibroblasts to activated, contractile myofibroblasts capable of enhanced extracellular matrix production and tissue remodeling.

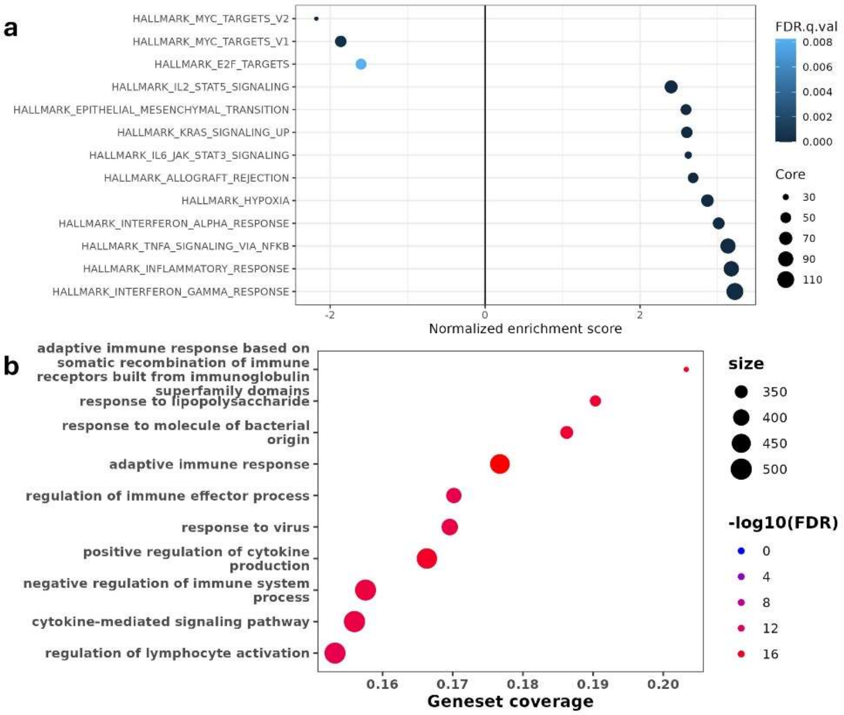


**Supplementary Figure 8: VAT EV treatment induces pro-inflammatory activation and immune response pathways in macrophages. a,** Hallmark pathway enrichment analysis showing normalized enrichment scores for significantly altered pathways in VAT EV-treated macrophages compared to controls. VAT EVs strongly upregulate inflammatory pathways including interferon gamma and alpha responses, TNF-α signaling via NF-κB, and general inflammatory response, consistent with M1 macrophage polarization. Additional upregulated pathways include IL2-STAT5 and IL6-JAK-STAT3 signaling, epithelial-mesenchymal transition, KRAS signaling, allograft rejection, and hypoxia response, indicating comprehensive immune activation and stress response. Proliferative pathways (MYC targets, E2F targets) show negative enrichment, suggesting reduced proliferation in favor of inflammatory activation. **b,** Gene Ontology biological process enrichment highlighting immune-related processes including adaptive immune response pathways, response to lipopolysaccharide and bacterial molecules, regulation of immune effector processes, cytokine-mediated signaling, and lymphocyte activation. The enrichment of adaptive immune response, cytokine production regulation, and pathogen recognition pathways demonstrates that VAT EVs promote a comprehensive pro-inflammatory M1 macrophage phenotype with enhanced capacity for immune surveillance and inflammatory cytokine secretion.

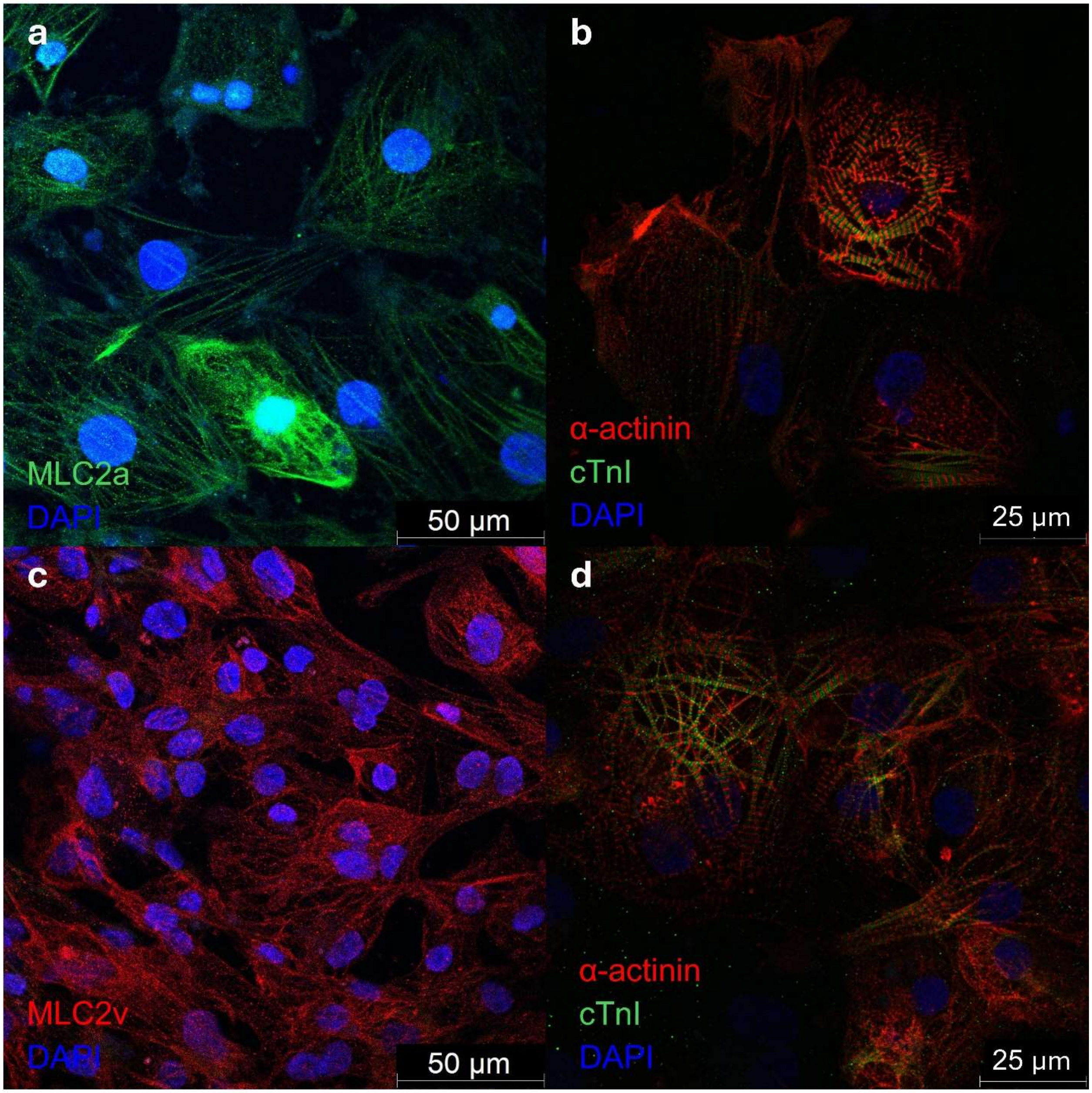


**Supplementary Figure 9: Characterization of iPSC-aCMs and iPSC-vCM. a-b,** Immunostaining of iPSC-aCMs shows expression of **a,** myosin regulatory light chain 2, atrial isoform (MLC2a or MYL7), **b,** sarcomeric α-actinin (α-actinin) and cardiac troponin I (cTnI). **c-d,** Immunostaining of iPSC-vCMs shows expression of **c,** myosin regulatory light chain 2, ventricular isoform (MLC2v or MYL2), **d,** sarcomeric α-actinin (α-actinin) and cardiac troponin I (cTnI).

**Supplementary Figure 10:**

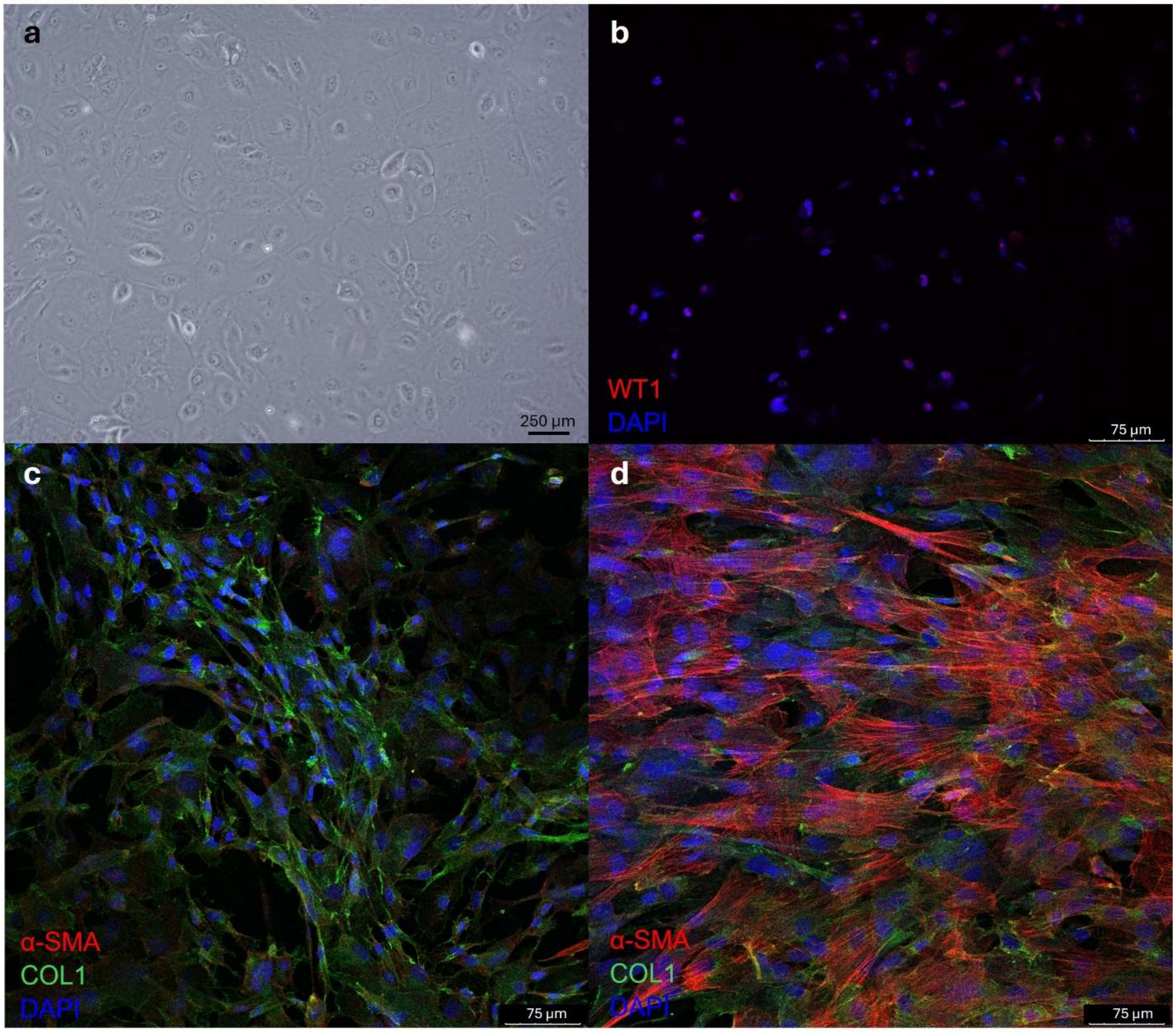


**Supplementary Figure 11: Characterization of iPSC-CFs. a,** Bright field image of epicardial cells, pre-cursor to cardiac fibroblasts, shows characteristic cobble stone appearance. **b,** Immunostaining of epicardial cells shows expression of WT1, epicardial cell maker, colocalizing with DAPT nuclear staining. **c,** Immunostaining of iPSC-CFs at resting state shows expression of collagen type 1 (COL1) with minimal expression of α-smooth muscle actin (α-SMA). **d,** Upon stimulation with transforming growth factor-β (TGF-β) at 10 ng/mL for 48 hours, there is significant increased expression of α-SMA, a marker of myofibroblasts.
